# Fitness cost associated with cell phenotypic switching drives population diversification dynamics and controllability

**DOI:** 10.1101/2023.04.06.535654

**Authors:** Lucas Henrion, Juan Andres Martinez, Vincent Vandenbroucke, Mathéo Delvenne, Samuel Telek, Andrew Zicler, Alexander Grünberger, Frank Delvigne

## Abstract

Isogenic cell populations can cope with stress conditions by switching to alternative phenotypes. Even if it can lead to increased fitness in a natural context, this feature is typically unwanted for a range of applications (e.g., bioproduction, synthetic biology, biomedicine…) where it tends to decrease the controllability of the cellular response. However, little is known about the diversification profiles that can be adopted by a cell population. We characterized the diversification dynamics for various systems (bacteria and yeast) and for different phenotypes (utilization of alternative carbon sources, general stress response and more complex development patterns). Interestingly, our results suggest that the diversification dynamics and the fitness cost associated with cell switching are coupled. For quantifying the contribution of the switching cost on population dynamics, we built a stochastic model that allowed us to reproduce the dynamics observed experimentally and identified three diversification regimes, i.e., constrained (at low switching cost), dispersed (at medium and high switching cost), and bursty (for very high switching cost). Furthermore, we used a cell-machine interface that we call the Segregostat to demonstrate that different levels of control can be applied to these diversification regimes, enabling applications involving more precise cellular responses.

## 1. Introduction

Cell populations can respond to environmental changes, and to the frequency of these changes, by adjusting phenotypes resulting from the activation of dedicated gene circuits^1–3^. This phenotypic plasticity has a lot of importance in microbial ecology, where the fitness of a cell population depends on a cost-benefit ratio between the sensing machinery needed for its activation and deactivation^4^ and the activity of a given gene circuit. Thus, controlling the phenotype of cells has a lot of importance in various fields of research, such as bioproduction and synthetic biology, where coordinated gene expression is typically wanted^5–10^. Generating and controlling cell collective behavior is considered as a hallmark of synthetic biology^9,11,12^, and is now enabled by the parallel advances made at the level of cell cultivation procedures (i.e., microfluidics^13^ and cell-machine interfaces^14^), as well as the manipulation of synthetic gene circuits^15–17^. Effective control of gene expressions and their underlying cellular functions can be achieved in cell populations^5,18,19^ or individual cells within a population^20,21^. Different approaches can be used to coordinate/synchronize gene expression in cell populations. On the one hand, specific gene circuits can be designed in order to generate natural oscillations^12,22^. On the other hand, external forcing can be used for coordinating cellular responses^6,23,24^. According to this last approach, a given stimulus (e.g., chemical inducer^20^, light^18,25^…) is repeatedly applied at a given frequency and amplitude in order to entrain gene expression within a cell population. In this case, the effective transfer of information from the extracellular environment to the effector sites within cellular systems is of critical importance and can be corrupted by biological noise^26–28^. In silico experiments pointed out that specific environmental fluctuation frequencies could significantly reduce stochasticity in cell switching, giving rise to corresponding population diversification regimes^1,28^. However, the main factors affecting these diversification regimes are not known. This feature will be experimentally investigated in this work by looking at the temporal diversification profile of different types of cell populations in continuous culture. For this purpose, chemostat runs will be complemented by experiments conducted in Segregostat. The Segregostat relies on a cell-machine interface to generate environmental perturbations that are compatible with the diversification rate of the considered cell population (**Figure 1D**)^19^. This “rational” environmental forcing allows for the observation of several diversification cycles in one experimental run (**Figure 1E**). We applied this technology to look at the dynamics of cell populations with cellular functions leading to different fitness costs, i.e., utilization of alternative carbon sources (*E. coli*), general stress response (*E. coli* and *S. cerevisiae*), sporulation (*B. subtilis*) and activation of a T7-based expression system (*E. coli*). We determined that, based on the fitness cost associated to the cell switching mechanism (referred as switching cost or fitness cost in this study), three different population diversification regimes, with different level of sensitivity to environmental perturbations, can be observed.

**Figure 1:**
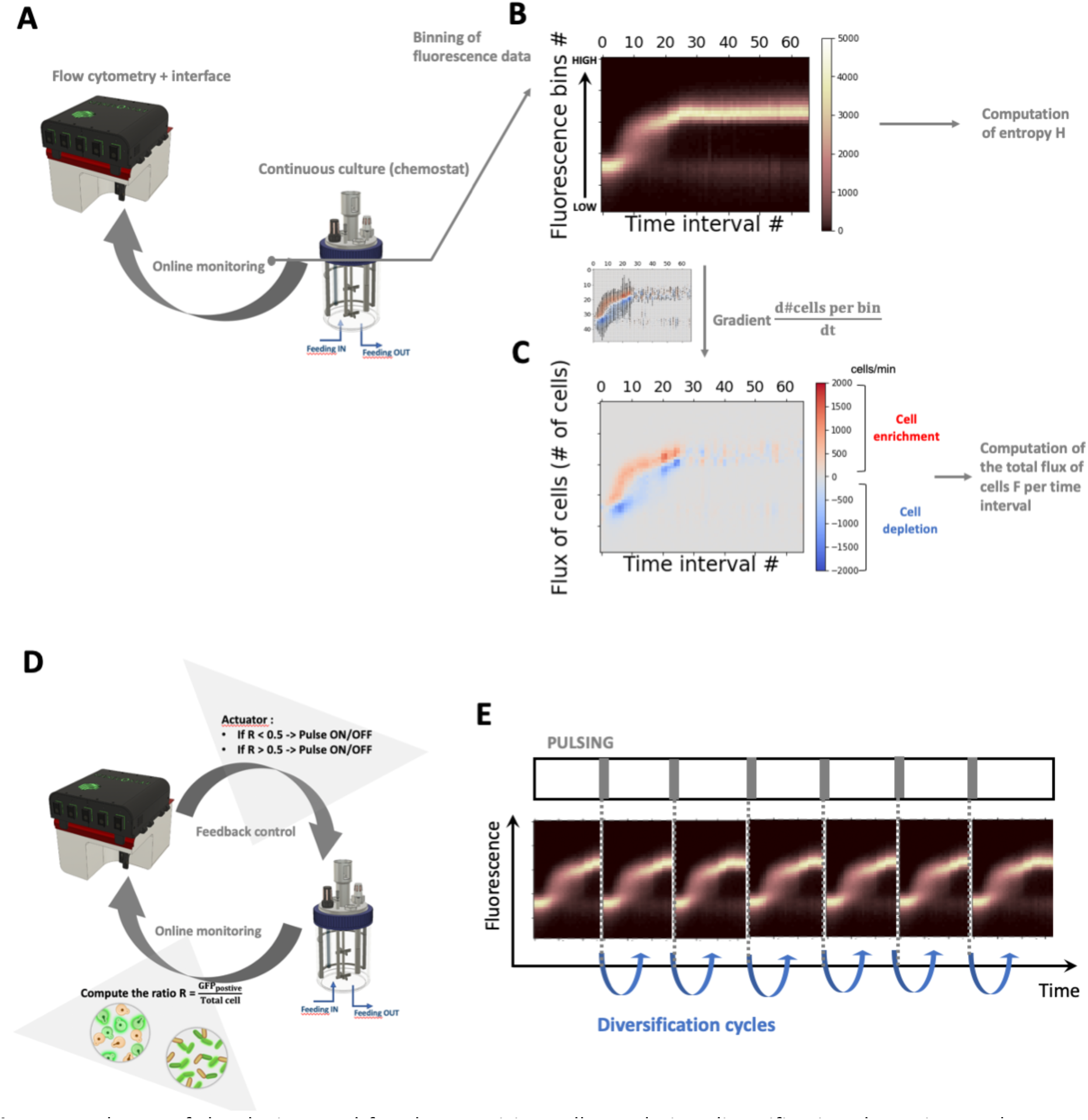
scheme of the device used for characterizing cell population diversification dynamics. **A** Chemostat culture is monitored based on automated FC. **B** The fluorescence distribution acquired by FC are assembled into a time scatter plot. This time scatter plot is then further reordered into 45 fluorescence bins in order to compute the evolution of the entropy H of the population (**Supplementary File, Supplementary note 1**). **C** The binned data are further processed by applying a gradient in order to compute the fluxes of cells into the phenotypic space, leading to the quantification of the total fluxes of cells per time interval F. **D** Scheme of the Segregostat set-up. Pulses of nutrient are added in function of the ratio between GFP negative and GFP positive cells, as recorded by automated FC. **E** Expected evolution of a Segregostat experiment where, upon controlled environmental forcing, several diversification cycles can be generated.

## 2. Results

### 2.1. Environmental forcing triggered by cell switching dynamics leads to coordinated gene expression for diverse biological systems

Temporal diversification of cell populations has been followed based on chemostat cultivation of GFP reporter bearing strains and automated flow cytometry (FC) (**Figure 1A**). In order to characterize population dynamics based on snapshot data, we developed a methodology to compute the fluxes of cells from a phenotype to another and the resulting degree of heterogeneity of the population, i.e., in our case, based on the measurement of information entropy (**Figure 1B&C**). This entropy is a measure derived from information theory allowing to compute the degree of heterogeneity of a population^29^. Briefly, GFP distributions obtained from automated FC measurements are binned and these bins are used to compute the cumulative probabilities of occurrence (**Supplementary file, Supplementary note 1, figure S1**). The benefit of this proxy is its independent from the mean of the distribution, by contrast with other noise proxies (e.g., Fano factor) that are known to be overestimated when the mean value increases^30,31^. Using entropy to analyze chemostat experiments, however, provides limited information about diversification dynamics. The main diversification process taking place during the transition between the batch and continuous phases of the culture. Therefore, we used a cell-machine interface allowing to produce several diversification cycles in a single experiment. This device is called the Segregostat and comprises a continuous cultivation device connected to an in-house online flow cytometry (FC) platform^19^ (**Figure 1D**). This device lets us observe several diversification cycles per experiment, leading to a better characterization of the population switching dynamics (**Figure 1E**). Practically, the cells analyzed based on automated FC are clustered into a GFP negative and a GFP positive group. Depending on the gene circuits used, a pulse of inducer is applied when the ratio between the two clusters (i.e., either 50% or 10% of the total amount of cells in the desired state, depending on the cellular system considered) is not met.

This methodology was applied to map the diversification of cell populations upon chemostat and Segregostat cultivations. We began our analysis by considering two gene circuits involved in simple cellular processes in *E. coli*, i.e., the activation of the arabinose (**Figure 2A**) and lactose (**Figure 2B**) operons respectively. This type of cellular process is quite simple since it involves two inputs, i.e., the absence or limitation in glucose and the presence of either lactose or arabinose as an alternative carbon source^4^. Since these cultures were carried out in continuous mode, it was quite easy to ensure glucose limitation. Furthermore, the gene circuitry behind the activation/deactivation of these two operons is well documented in the literature^4,32^.

**Figure 2:**
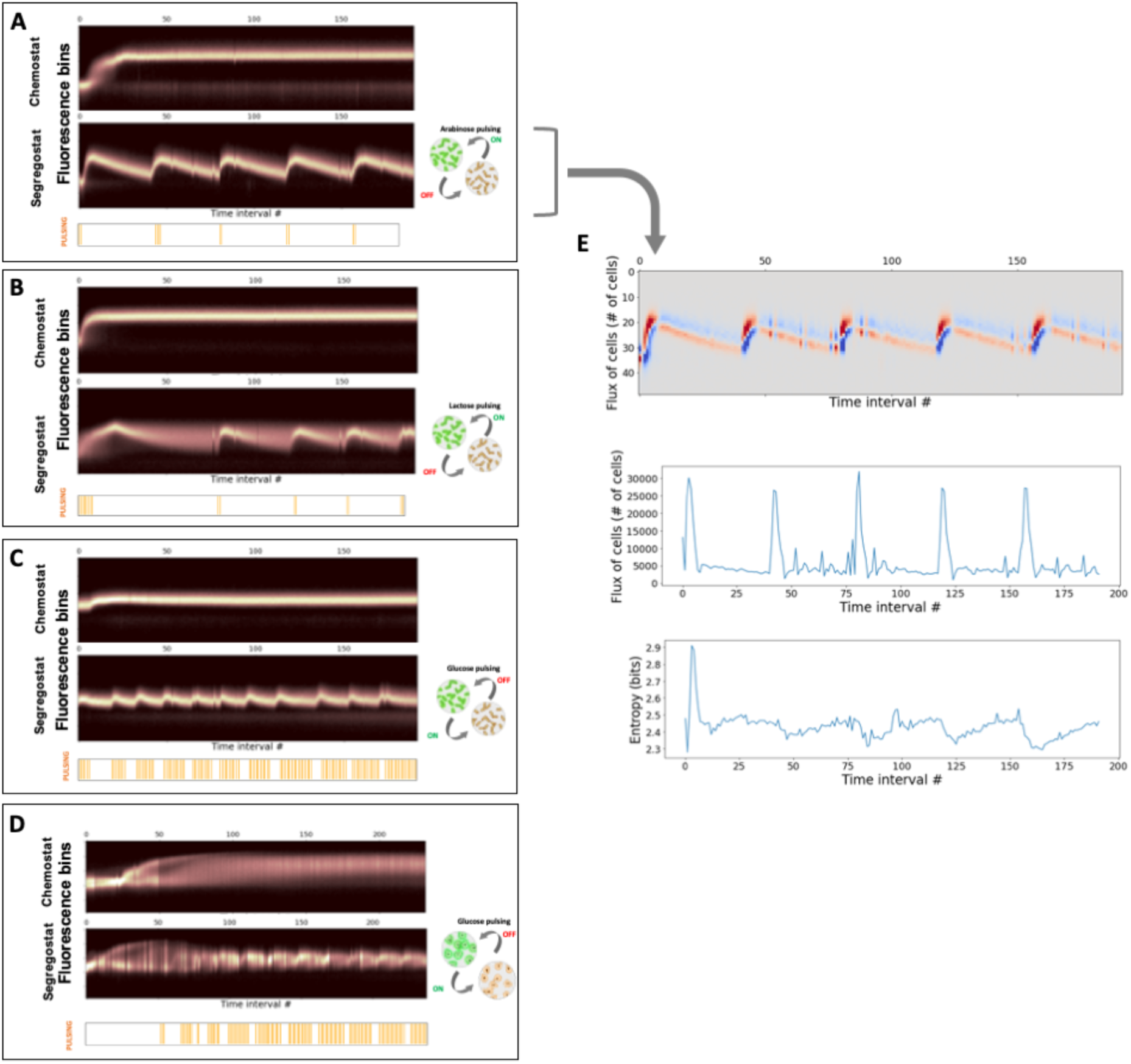
time scatter plots binned into 50 cell clusters (fluorescence bins) for cultivations made in chemostat and Segregostat for **A** the P_*araB*_:GFP system in *E. coli*. **B** the P_*lacZ*_:GFP system in *E. coli*. **C** the P_*bolA*_:GFP system in E.coli. **D** the P_*glc3*_:GFP system in *S. cerevisiae*. Animated movies for the time evolution of the FC raw data for each system are available (**Supplementary Movies M1** to **M8**). For the Segregostat experiments, environmental forcing has been performed based on nutrient pulsing (type of nutrient shown in the drawings for cell switching). **E** Computation of the flux of cells and the entropy for the P_*araB*_:GFP system cultivated in Segregostat mode (similar computation have been done for the other systems and can be found in **Supplementary information**, **Figure S2**). All the data are displayed in function of time intervals of 12 minutes, as the population snapshots have been acquired by automated FC.

A feature of these two systems is that, upon activation, these circuits do not result in a reduction of fitness. More often than not, that is not the case in natural gene circuits. We thus decided to investigate other gene circuits involved in more complex cellular processes, and known to lead to a higher switching cost (growth reduction). We first chose to consider the general stress response in *E. coli* and selected the promoter of the *bolA* gene as a representative σ^S^-dependent system^33,34^ (**Figure 2C**). In order to extend our analysis to another biological system, we also selected a promoter involved in the accumulation of glycogen in yeast^35^, i.e., P_*glc3*_, as a representative reporter of bet-hedging in *S. cerevisiae*^36,37^ (**Figure 2D**). Both genes are involved in very complex regulons, making it difficult to find an external trigger. However, both reporter systems are known to share common features in the sense that the growth of individual cells is anticorrelated with the level of expression of these general stress response reporters, making them very useful for analyzing cell collective behavior such as bet-hedging^34,38^. We then decided to use the external glucose concentration as the main actuator for these two systems. Glucose-limited chemostats were then run as reference conditions. For the Segregostat experiments, glucose was pulsed instead of lactose or arabinose, allowing to generate feast to famine environmental transitions.

For all four cellular systems investigated, Segregostat cultivation led to entrainment and sustained oscillation of gene expression (**Figure 2A-D**). Based on the analysis of the entropy H(t) and the flux of cells F(t) over time, we observed that for all systems, entrainment phases were accompanied by an increased flux of cells switching to the alternative phenotype and a corresponding decrease of entropy H(t) at the time of pulsing (**Figure 2E** where the analysis is shown for the P_*araB*_:GFP system; analyses for the other systems can be found in **Supplementary file, Supplementary note 1, figure S2**). However, the entropy increases during the relaxation phase (GFP dilution upon cell division). At this stage, the question is whether the different population profiles recorded based on automated FC can lead to different functionalities. We then decided to conduct an in-depth characterization of the heterogeneity of cell populations.

### 2.2. Coordinated gene expression doesn’t necessarily lead to a more homogeneous cell population

We then compared the entropies recorded for each systems considered cultivated either in chemostat or Segregostat modes (**Figure 3A-D**). For the P_*araB*_:GFP and P_*bolA*_:GFP only slight differences of H were observed between chemostat and Segregostat conditions. However, for the two other systems, noticeable differences were observed. For the P_*lacZ*_:GFP, the average entropy in the Segregostat is always higher than in chemostat. This is probably due to the leakiness of the promoter during the relaxation phase of the diversification cycles. This causes unwanted induction even when little to no inducer is present. On the other hand, we observed a significant reduction in entropy for the P_*glc3*_:GFP system when cultivated in Segregostat, suggesting that cell entrainment can have an effect on the structure of the population. In order to understand why we do observe such differences in behavior between the cellular systems, we computed the mutual information (MI) between the environmental conditions and the activation of the target gene circuit for the P_*araB*_:GFP and P_*glc3*_:GFP systems. MI is a proxy derived from information theory^26,27^ and involves the computation of the entropy of the cell population, as defined in the previous section. In short, MI tells us how much we can learn about the input (i.e., in our case the environmental stimulus used for entraining the cell population) from the output (i.e., in our case the distribution of GFP in cell population, the dispersion being quantified based on the entropy). Thus, in our case, MI is a proxi for information transfer efficiency between the inducer concentration and the cell population induction. The total entropy for each system was evaluated from the conditional probabilities obtained by exposing cell populations to different cultivation conditions (**Figure 3E&F**). Detailed description of the experiments done for determining the conditional probability distributions can be found in **Supplementary file, Supplementary note 2, Figures S3–S6**. MI was obtained by subtracting the time-dependent entropies to the total entropies recorded for each system. For the P_*araB*_:GFP system, MI is already relatively high in the chemostat leaving little room for improvement in the Segregostat (**Figure 3C&E**). It means that a reduction in entropy between the two cultivation modes has to be expected when the amount of information conveyed in chemostat condition is low (e.g., when cells cultivated in chemostat do not sense the inducer and, accordingly, do not activate the corresponding gene circuit). This is exactly what happened during the chemostat culture of the P_*glc3*_:GFP system (**Figure 3D**). MI analysis had pointed out that there was still room for additional reduction in entropy (**Figure 3D&F**), and this was observed in Segregostat where glucose pulses reduced the average entropy (**Figure 3D**). This indicates that P_*glc3*_:GFP is the only studied system where entrainment leads to a more homogeneous population. The next section investigates why such a phenomenon occurs.

**Figure 3:**
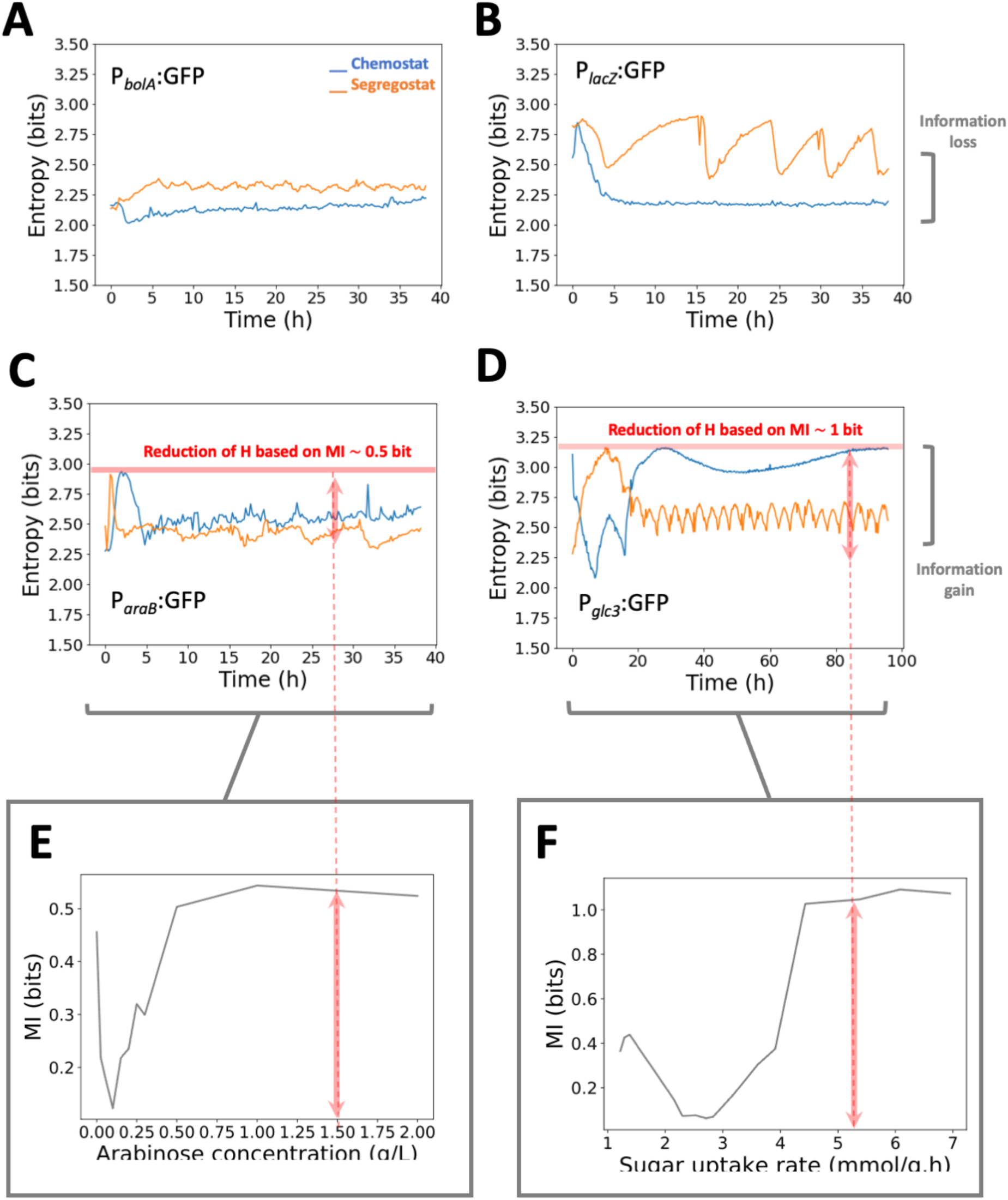
Comparative analysis of the entropy computed from the binned population data for four different cell systems. Comparative analysis of the entropy profile for the **A** P_*bolA*_:GFP system in *E. coli*. **B** P_*lacZ*_:GFP system in *E. coli*. **C** P_*araB*_:GFP system in *E. coli*. **D** P_*glc3*_:GFP system in *S. cerevisiae*. For the P_*araB*_:GFP and P_*glc3*_:GFP systems, the conditional probabilities, i.e., the GFP distribution of the population exposed at different environmental conditions, have been experimentally determined (**Supplementary information**, **Supplementary note 2**), allowing the computation of the mutual information (MI). **E** MI for the P_*araB*_:GFP system exposed to different arabinose concentrations. The MI distribution for the P_*araB*_:GFP system suggests that, at high arabinose concentration a gain of information of approximately 0.5 bit has to be expected (the value has been reported by a red line on **Figure 3C**). **F** MI for the P_*glc3*_:GFP system exposed to different sugar uptake rates in a accelerostat cultivation device (**Supplementary information**, **Figure S4**). The MI distribution for the P_*glc3*_:GFP system suggests that, at high glucose concentration a gain of information of approximately 1 bit has to be expected (the value has been reported by a red line on **Figure 3D**).

### 2.3. Fitness cost associated to cell phenotypic switching leads to a higher entropy at the population level that can be reduced upon environmental forcing

Stress response pathways in yeast are known to be involved in bet-hedging strategies, leading to a trade-off between growth and expression of stress-related genes^37,38^. The *glc3* gene belongs to this category. Accordingly, cells activating *glc3* exhibit reduced growth. This phenomenon has been characterized based on Microfluidics Single-Cell Cultivation (MSCC)^39^ experiments allowing to expose yeast cells to tightly defined glucose concentrations (**Figure 4A**). Unlike with FC analyses where only population snapshots are captured, these experiments let us monitor cell traces and thus analyze the fitness cost (growth reduction upon switching) associated to the switching. It can be observed that, at low glucose concentration (< 0.2 mM), a single cell fully activates stress reporter and stops growing (**Supplementary movie M12**). At a higher glucose concentration, growth of the microcolonies is faster, but some stochastic switching events can be clearly observed, with cells suddenly expressing the fluorescence reporter and stopping their growth (**Supplementary movie M13**). In order to confirm the beneficial impact of Segregostat condition on the P_*glc3*_:GFP, we used dynamic microfluidic single-cell cultivation (dMSCC)^40^ where we applied environmental fluctuations between 0.1 and 1 mM of glucose, at the frequency recorded in the Segregostat conditions (**Figure 2D**). We observed a very homogenous gene expression pattern with cells turning green in perfect synchrony (**Figure 4B, Supplementary movie M14**). The growth of all cells was comparable, the stress level being kept at a low level thanks to the fluctuating environmental conditions. It seems that when phenotypic switching is associated with a loss of fitness, there is more stochasticity in the diversification pattern followed by the population. However, when the nutrient level is changed at a given frequency, switching can be kept under control, leading to a drastic reduction of the phenotypic heterogeneity of the cell population.

**Figure 4:**
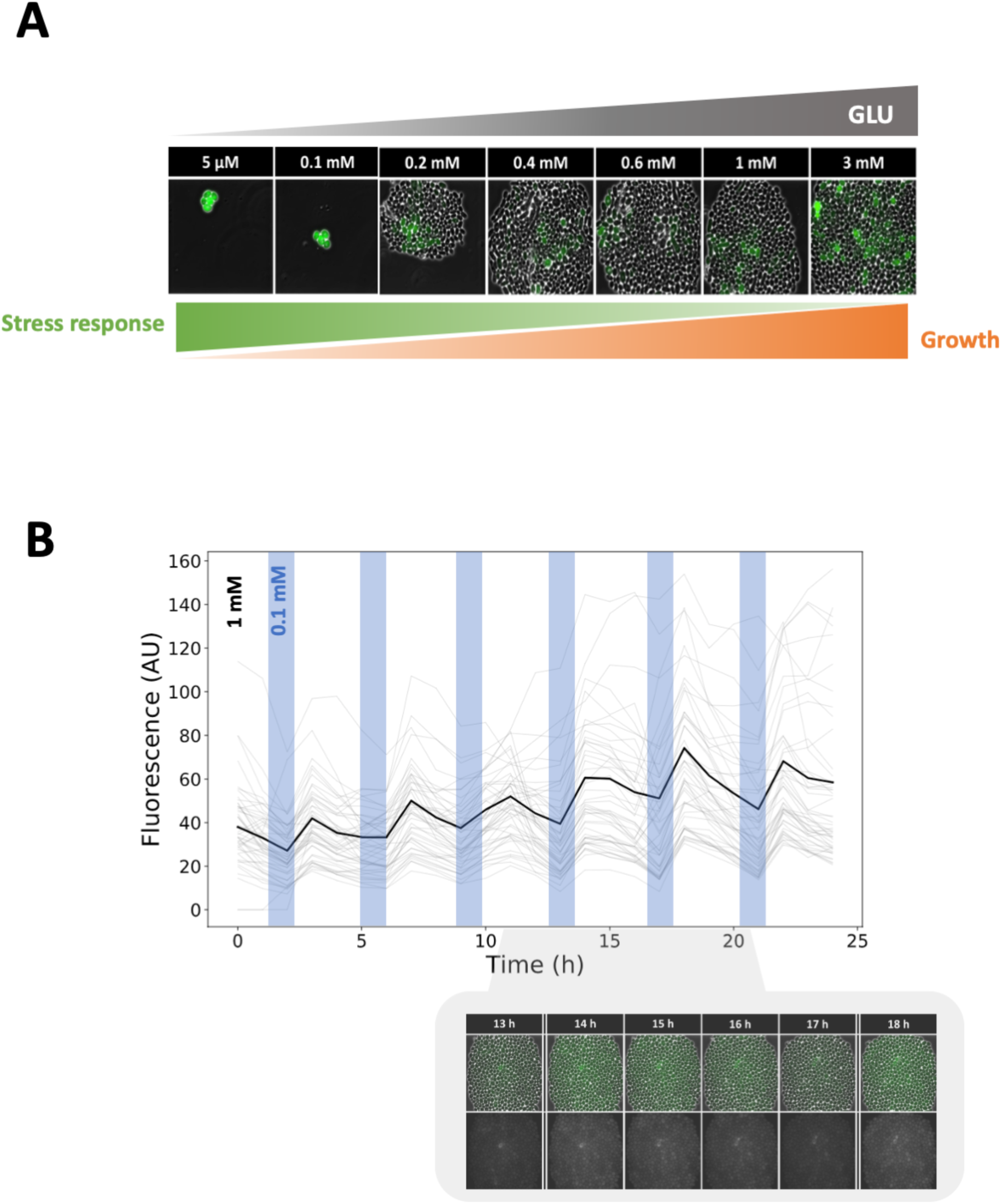
**A** Picture of yeast P_*glc3*_:GFP microcolonies cultivated in a MSCC device at different glucose concentrations. Movies of a microcolony growing at a concentration of 0.1 mM and 1mM can be accessed in **Supplementary movie M12** and **M13** respectively). **B** Single cell traces of yeast P_*glc3*_:GFP cells cultivated in a dMSCC device fluctuating between 1 and 0.1 mM of glucose (T_1mM_ = 3h; T_0.1mM_ = 0.8h). Between 10 and 40 cells have been tracked in four different cultivation chambers over two biological replicates (mean fluorescence is shown in bold). Pictures of a microcolony taken at regular time intervals are shown (**Supplementary movie M14**).

### 2.4. Phenotypic switching associated with extreme fitness cost gives rise to a bursty diversification regime

According to the results exposed in the previous section, a lower entropy in Segregostat has to be expected when the fitness cost associated with phenotypic switching is high. To validate this hypothesis, we investigated the diversification dynamics of two other systems known for their high impact on cellular fitness; the T7-based expression system in *E. coli* BL21 – a typical heterologous protein production platform – and the sporulation regulon in *B. subtilis* (P_*spoIIE*_:GFP). Indeed, for both, phenotypic switching leads to a drastic reduction of growth. Surprisingly, even in chemostat, FC profiles reveal bursts of diversification (**Figure 5A&B**). These bursts are the result of a subpopulation of cells deciding to switch and being washed out from the continuous cultivation device due to the associated fitness cost. During chemostat cultivation, bursts involving small fluxes of cells occur continuously and result in a high entropy at the population level. Then, upon environmental forcing based on Segregostat cultivation, the number of bursts is reduced and the fluxes of cells involved in the process are increased, leading to a substantial but temporary reduction of the entropy for the population (**Figure 5A&B**).

**Figure 5:**
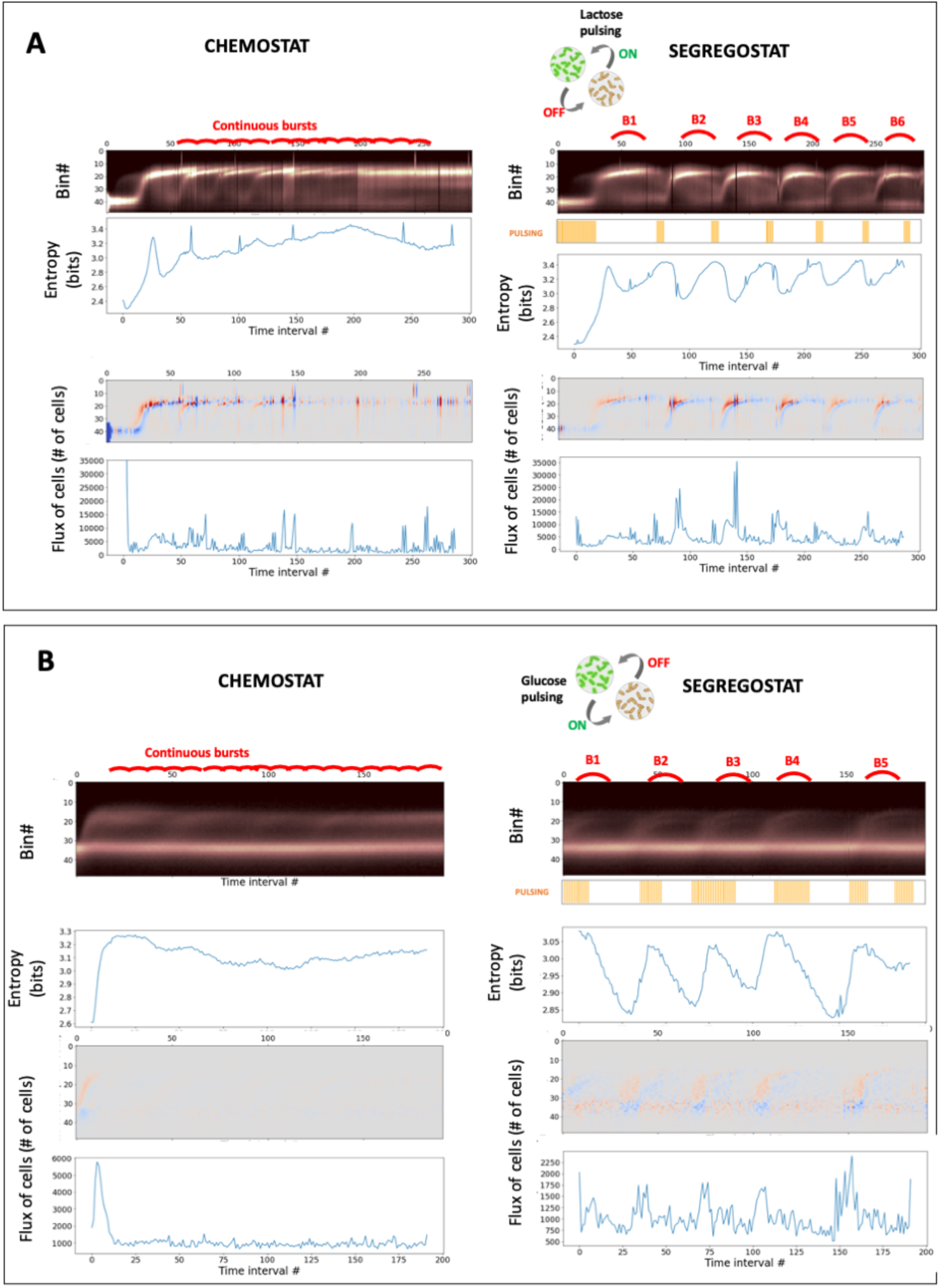
Temporal diversification profile for the P_*T7*_:GFP system in *E. coli* (**A**) and the P_*spoIIE*_:GFP system (**B**) cultivated in chemostat and Segregostat modes. Only the continuous phase of the cultivation is shown on the graphs. Bursts of diversification are highlighted in red on the fluorescence bins data (the full diversification profile determined based on automated FC is available as **Supplementary movie M9-M10**. In both cases, the entropy H(t) and the fluxes of cells in the phenotypic space F(t) have been computed from the binned fluorescence data. All the data are displayed in function of time intervals, as the population snapshots have been acquitted by automated FC. One time interval corresponds to 12 minutes.

These results point out that the Segregostat reduces the average entropy of gene circuits with a high fitness cost despite very complex dynamics. It has been suggested in the literature that the stochasticity in cell switching is associated to its fitness cost and is important for the survival of the whole population. This feature is well illustrated in this case where a phenotype switch induces a dramatic loss of growth rate, leading to the wash-out of these cells in continuous cultivation conditions. However, this stochasticity can be reduced by applying environmental perturbations at a rate matching the phenotypic switching rate of cells. In the context of the T7 system, this approach led to the maximization of cells in the GFP positive state suggesting that it could be used for mitigating metabolic burden and maximizing productivity in continuous bioprocesses.

### 2.5. Fitness cost drives the appearance of different dynamical regimes with different levels of controllability

According to the datasets acquired from the different biological systems, we observed three types of diversification regimes named respectively constrained, dispersed and bursty (**Figure 6**). When the fitness cost associated to the switching is low or non-existent, we observe a constrained diversification regime where the population switches upon environmental change and adopts a homogeneous distribution. In that case, the stimulation of the population with controlled environmental pulsing does not homogenize it (since information transmission is already maximal in the non-controlled conditions, **Figure 3A&B**).

**Figure 6:**
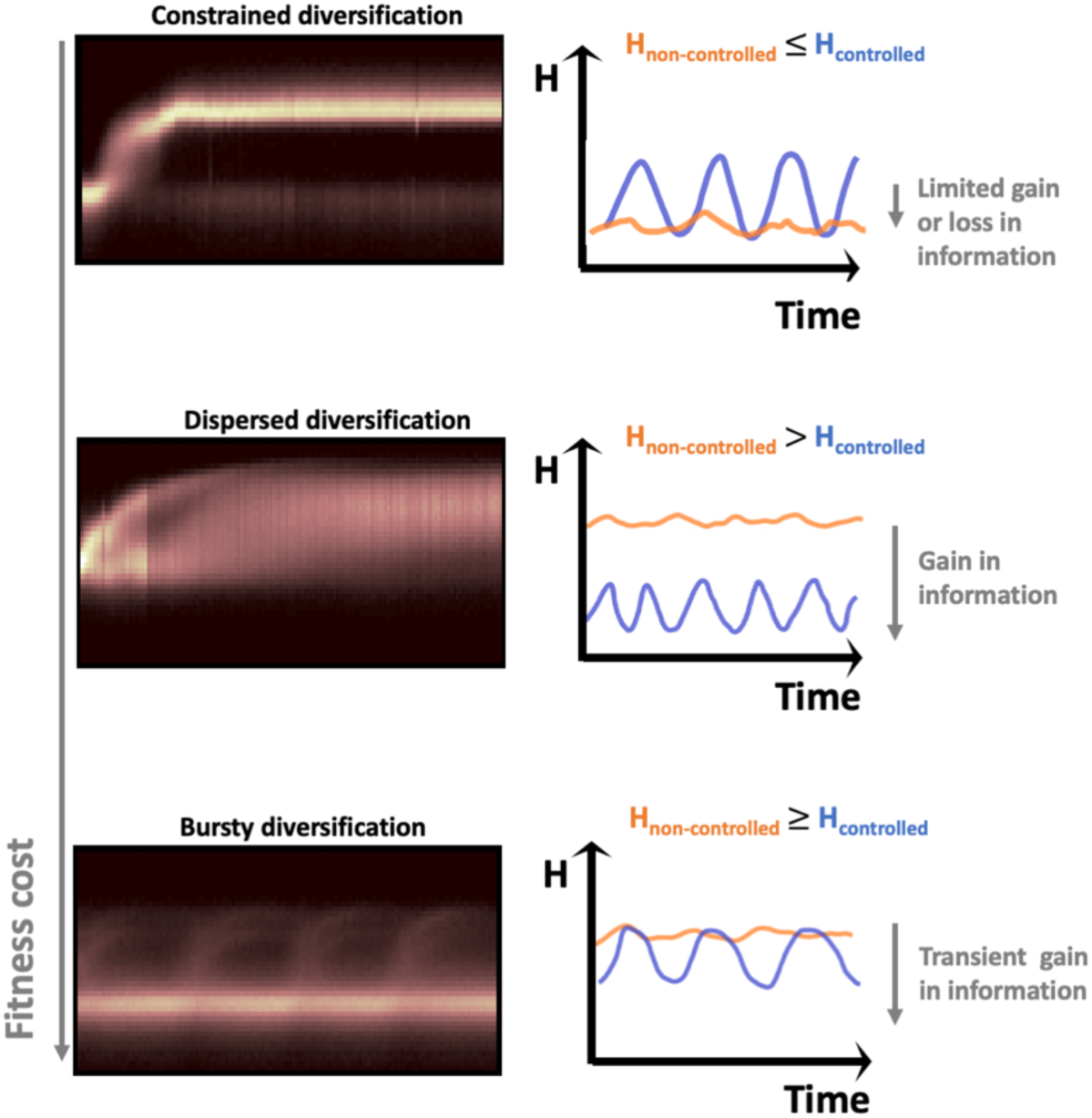
illustration of the three diversification regimes observed based on automated FC in function of the fitness cost. The consequences on the control of cell population are also shown based on the time evolution of the entropy of the population between non-controlled (i.e., obtained upon standard chemostat cultivation) and controlled (i.e., obtained upon periodic environmental stimulation of the population in the Segregostat) conditions.

When there is a fitness cost associated to the switch, what we call a dispersed diversification regime can be observed. In that case, cells react to environmental changes but then adopt a more heterogeneous population structure. In this case, the application of controlled environmental perturbations allowed a substantial reduction in population heterogeneity. For the bursty diversification regime (higher fitness cost), cells switch in bursts, leading to a very heterogenous population structure. The application of controlled environmental perturbations reduces the number of bursts and increases the number of cells involved in these bursts, leading to a transient decrease in population heterogeneity.

It seems that the regime depends on the fitness cost associated with the phenotypic switching event. In order to verify that fitness cost is indeed the driver for the diversification pattern adopted by cell populations, we conducted in silico experiments. For this purpose, we considered the kinetic parameters obtained from the inference of the P_*glc3*_:GFP system in yeast and conducted stochastic simulations based FlowStocKS (**Supplementary file, Supplementary note 3**). We conducted 32 different chemostat simulations by varying only the value for the fitness cost and computed the entropy H (**Figure 7A**) and the fluxes of cells (F) involved in phenotypic switching (**Figure 7B**). Solely based on the fitness cost associated to the switching, we were able to reproduce the three types of diversification regimes experimentally observed during the experiments (**Figure 3C**). Complete wash-out of the cells was observed for extreme fitness cost (>99% reduction in growth rate). We then wondered if we could observe clear transitions between the different regimes. Such transition was observed between the bursty and the dispersed regime based on the computation of the flux of cells F. Indeed, the bursty regime is marked by the appearance of a strong variation in flux of cell which is not observed for the other two regimes (**Figure 7E**). The transition between the dispersed and constrained regime is more progressive (**Figure 7D**). FlowStocKS was also able to reproduce the behavior of the population under Segregostat cultivation, and the reduction in entropy upon environmental forcing was computed (**Figure 7F**). Again, reduction in entropy depended on the associated fitness cost and thus was observed for the dispersed and bursty regimes, in accordance with our experimental observations.

**Figure 7:**
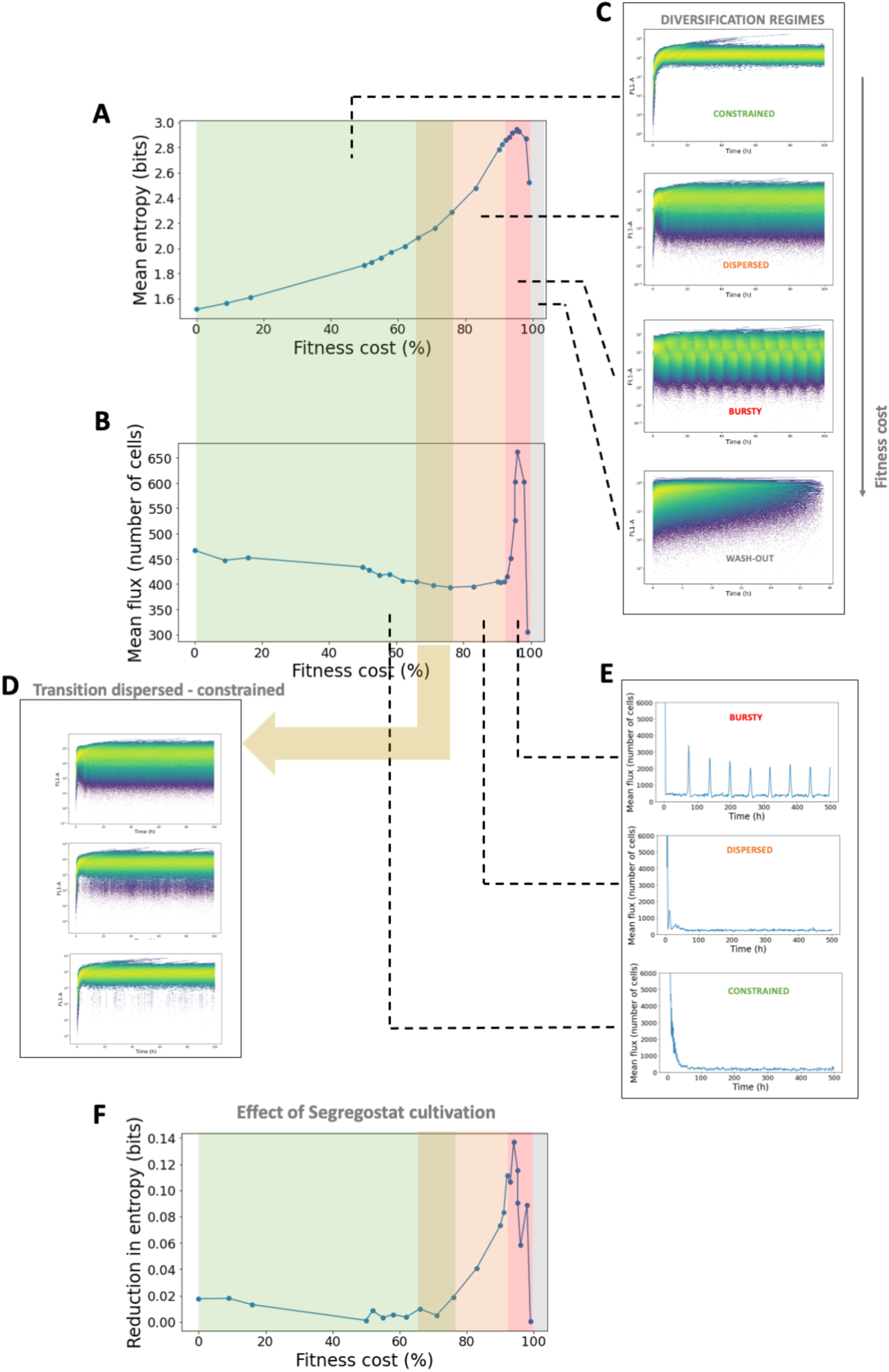
main outputs of the FlowStocKS simulations. **A** Evolution of the mean entropy recorded in chemostat in function of the fitness cost (here expression as the percentage of reduction of the initial growth rate prior phenotypic switching). **B** Evolution of the mean flux of cells recorded during chemostat experiments in function of the fitness cost associated to phenotypic switching (see **Figure 1** for more details about the computation of the flux of cells). **C** Selected simulated time scatter fluorescence plots illustrating the different diversification regimes observed at different fitness costs (the whole simulation dataset can be found as **Supplementary movie M15**). **D** Selected simulated time scatter fluorescence plots illustrating the progressive transition between the dispersed and constrained diversification regimes. **E** Selected simulated time evolution for the fluxes of cells recorded for different values of fitness cost. The bursty regime is characterized by the spontaneous generation of flux of cells (bursts) in chemostat cultivations. **F** Reduction of the entropy upon environmental forcing in function of the fitness cost associated with phenotypic switching. The entropy values have been computed by substracting the mean entropy value recorded for chemostat experiment to the corresponding ones obtained in Segregostat.

## 3. Discussion

We used the Segregotat to better characterize cell population diversification dynamics by generating successive diversification cycles during the same experimental run. The basic principle behind this technology is to revert the environmental conditions when a fraction of cells (50% or 10% of the total population depending on the system investigated) crossed a predefined fluorescence threshold. This approach allows to maintain a cell population in a dynamic switching state during the experiment. Based on the analysis of the mutual information, i.e., the amount of information transferred from the extracellular conditions to the cell systems^27,29,42^, we determined that for the P_*araB*_:GFP system, the chemostat drives a similar amount of information than the Segregostat. On the other hand, we observed a drastic reduction in entropy when entraining stress-related systems, such as the P_*glc3*_:GFP system in yeast, in the Segregostat. In this case, we determined that the high entropy observed in the chemostat was related to a trade-off between growth and gene expression^36–38,43–45^, which was further confirmed based on microfluidics experiment, and by considering two additional systems where phenotypic switching induced a huge fitness cost i.e., the sporulation system in *B. subtilis* and the T7-based expression system in *E. coli*.

Based on all the data accumulated by automated FC for six different biological systems, we found that cell populations diversified according to three distinct regimes associated to increasing fitness costs, i.e., constrained, dispersed and bursty. The most noticeable difference between these regimes is the level of entropy of the cell population, the entropy being a measure derived from information theory giving a robust estimate of population dispersion^29,41^. The lowest entropy values were observed for the constrained regime and the highest ones were observed for dispersed and bursty regimes. The other difference was observed upon the cultivation of cell populations under fluctuating environmental conditions in the Segregostat were a reduction of entropy compared to chemostat cultivation was observed for the dispersed and bursty regimes but not the constrained. Taken altogether, the data suggested that cell population diversification dynamics are mainly driven by the fitness cost associated with the phenotypic switching mechanism, a higher cost giving rise to the dispersed and bursty regimes. All these observations were confirmed based on stochastic simulations (FlowStocKS), suggesting that the proposed diversification framework could be generalized for characterizing diversification dynamics for any kind of cellular system.

Harnessing phenotypic heterogeneity of microbial populations has been the subject of much research, leading to the design of various technologies aiming at homogenizing gene expression in cell populations^46^. We have shown that the level of diversification of microbial populations cultivated in continuous bioreactors depends mainly on the fitness cost. Since many applications involve gene circuits whose activation leads to substantial burden for the cellular system^48,49^, active diversification processes have to be expected in a number of cases^50^. Our data point out that, under these circumstances, phenotypic diversity could be promoted, instead of being reduced, by using environmental forcing to provide cell populations with robust temporal patterns in gene expression. For example, bursty diversification profiles have been observed for two cellular systems exhibiting high switching cost. According to this regime, marked cycles of diversification can be observed even in chemostat cultures. These cycles are due to the rapid switching (burst) of a fraction of the population that lower the average growth rate of the population and are washed out of the system. These cells are then replaced by the next burst of diversification, starting from a subpopulation of non-diversified cells. This type of temporal profile has been previously observed, but based on spatially organized cells equipped with synthetic circuits^7,51,52^. Here, we show that it is possible to reproduce such a complex but organized diversification profile with cells in suspension in a bioreactor and that the complex dynamics behind phenotypic heterogeneity are linked to the fitness cost associated to the switch.

## 4. Methods

### 4.1. Stains and plasmids

The analyses of alternative carbon source utilization and stress response in *E. coli* were done based on a *E. coli* W3110 backbone and kanamycin resistance bearing plasmids originating from the Zaslaver collection (i.e., *P_araB_:*:GFPmut2, *P_lacZ_:*:GFPmut2 and *P_bolA_:*:GFPmut2)^53^. For investigating the T7 induction system response, we used *E. coli* BL21 (DE3) carrying pET28:GFP^56^. To observe the starvation response, we used *S. cerevisiae* CEN-PK 117D background with the chromosomal integration of a reporter cassette *P_glc3_:*eGFP^35,54^. The *E. coli* W3110 *ΔaraBAD* strain used for determining the conditional entropy of the P_*araB*_:GFP system was constructed through CRISPR-cas9 enhanced lambda red phage mediated homologous recombination as described by Jiang *et al.*^55^ (the primer sequences are described in **Supplementary File, Table S1**). Finally, we monitored the early stage of the sporulation process in *B. subtilis* 168 based on a chromosomal integration of *P_spoIIE_:*:GFPmut2 (kindly provided by Denise Wolf and Adam Arkin)^57^.

### 4.2. Cultivation conditions and Segregostat procedure

Bacteria (*E. coli* and *B. subtilis)* precultures and cultures have been performed in a defined mineral salt medium containing (in g/l): K_2_HPO_4_ 14.6; NaH_2_PO_4_.2H_2_O 3.6; Na_2_SO_4_ 2; (NH_4_)_2_SO_4_ 2.47; NH_4_Cl 0.5; (NH_4_)2-H-citrate 1; glucose 5, thiamine 0.01, antibiotic 0.1. Thiamine is sterilized by filtration (0.2 mg/l). The medium is supplemented with 3 ml/l of a trace element solution, 3ml/l of a FeCl_3_.6H_2_O (16.7 g/l), 3 ml/l of an EDTA (20.1 g/l) and 2ml/l of a MgSO_4_ solution (120 g/l). The trace element solution contains (in g/l): CoCl_2_.H_2_O 0.74; ZnSO_4_.7H_2_O 0.18; MnSO_4_.H_2_O 0.1; CuSO_4_.5H_2_O; CoSO_4_.7H_2_O. Filtered sterilized kanamycin (50 mg/l) was added for plasmid maintenance in *E. coli*. *S. cerevisiae* cultures and precultures have been performed based on a Verduyn mineral medium^49^. The precultures were performed in 1L baffled flask overnight either at 37 °C (bacteria) or 30°C (*S. cerevisiae*) at a shaking speed of 150 rpm and used to start the batch phase in a lab-scale stirred bioreactor (Biostat B-Twin, Sartorius) total volume: 2 l; working volume: 1 l at an initial OD_600_ of 0.5. Once the batch phase was over (typically after 5,8 and 15 h for *E. coli, B. subtilis*, *S. cerevisiae* respectively). The operating conditions used for the various cell systems are provided in **Supplementary information**, **Supplementary note 4**, Table S3.

Data was collected through online flow cytometry during the experiments, and in Segregostat experiments, the actuator was pulsed based on both the observed distribution and a pre-defined set-point. The Segregostat platform has been described earlier^10^. Briefly, every 12 minutes, a sample is automatically taken from the bioreactor, diluted, and analyzed in a flow cytometer (BD Accuri C6, BD Biosciences) with an FSC-H analysis threshold of 20,000 for bacteria and 80,000 for *S. cerevisia*e. During the chemostat experiment, glucose and, if applicable, an alternative carbon source was fed together, while in the Segregostat experiment, glucose was the sole carbon source in the feed. In Segregostat, a feedback control loop, which includes a custom MATLAB script based on FC data, activates a pump to pulse an actuator. For *S. cerevisiae*, *E. coli* stress response, and *B. subtilis* sporulation, the actuator pulses glucose, while for *E. coli* alternative carbon source utilization, the actuator pulses lactose and arabinose. For all systems except for the sporulation control, a control threshold of 50/50 was utilized. However, for the sporulation control, the regulation was triggered once the fluorescence threshold was exceeded by more than 20% of the cells. This decision was made based on the irreversibility of the sporulation process, which required earlier intervention.

### 4.3. Microfluidic cultivations and time lapse microscopy

All devices and parameters were the same as previsouly^59^, excepted few modifications. *S. cerevisiae* cells have been cultivated in the dynamic microfluidic single-cell cultivation (dMSCC) chips provided by Alexander Grünberger’s lab (reference 24W, chambers size : 80 µm x 80 µm x 850 nm)^40^. Their design enables the simultaneous use of 2 cultivation media. They are separated in three zones: 2 control zones, fed by either one or the other medium, and a switching zone fed in alternance by the two media. Diverse combinations of Verduyn medium with different glucose concentrations have been tested (i.e., 0.4 mM - 0.6 mM, 0.2 mM – 0.8 mM, 0.1 mM - 1 mM and 5 µM – 3 mM). To approach the Segregostat conditions, the duration of the switching zone feeding with the medium containing more (or less) glucose corresponds to the mean period with (or without) pulsing when population is controlled in the segregostat (i.e., 180 min (or 48 min)). High precision pressure pumps (line-up series, Fluigent, Le Kremlin-Bicêtre, France) were used to precisely control medium flow rate. The temperature was set at 30◦C. The chambers were inoculated with one or two cells by flushing the device with a cell suspension (OD600 between 0.4 and 0.5). At least 6 cultivation chambers were selected manually for each zone of the dMSCC chips. Microscopy images were acquired during 72 hours using a Nikon Eclipse Ti2-E inverted automated epifluorescence microscope (Nikon Eclipse Ti2-E, Nikon France, France) equipped with a DS-Qi2 camera (Nikon camera DSQi2, Nikon France, France), a 100× oil objective (CFI P-Apo DM Lambda 100× Oil (Ph3), Nikon France, France). The GFP-3035D cube (excitation filter: 472/30 nm, dichroic mirror: 495 nm, emission filter: 520/35 nm, Nikon France, Nikon) was used to measure GFP. The phase contrast images were recorded with an exposure time of 300 milliseconds and an illuminator’s intensity of 30%. The GFP images were recorded with an exposure time of 500 milliseconds and an illuminator’s intensity of 2% (SOLA SE II, Lumencor, USA). During the first 48 hours, GFP and phase contrast images were acquired every hour. During the 24 last hours, phase contrast images were acquired every 6 minutes and GFP images every hour (or every 20 minutes for one experiment with media 5 µM – 3 mM glucose). The optical parameters and the time-lapse were managed with the NIS-Elements Imaging Software (Nikon NIS Elements AR software package, Nikon France, France). The single-cell data have been computed for the 24 last hours of the time lapse for at least 3 chambers per condition (i.e., zone of the dMSCC device). The cell-segmentation of the images and the measure of single-cell mean GFP intensity were performed using either the Python GUI^60^ or the Matlab algorithm of Wood and Doncic^61^. This last was also used for cell tracking. In this case, seeds for segmentations have been corrected manually and at least 10 cells per microfluidic cultivation chamber (well segmented and tracked all the time lapse long) were selected manually.

### 4.4. Modelling cell population dynamics based on FlowStocKS

The aim of this simulation toolbox is to be able to represent with high fidelity the population snapshots and dynamics captured based on automated flow cytometry. Briefly, population dynamics is modelled based on a set of ODEs representing the time evolution of biomass and substrates according to a Monod kinetics in a continuous cultivation device. From the global population, a given number of cells (approx. 10,000, some cells being washed-out during the simulation) are considered for generating a stochastic process. For these cells, phenotypic switching is modelled according to a Markov chain process driving the synthesis and degradation of GFP. Cell growth and division are taken into account for computing the GFP content. Upon switching, cells may encounter a fitness cost depending on an inhibitory kinetics. The data are then fitted to a seven-decade fluorescence scale in order to fit to the automated FC data. Detailed information, including parameters and equations settings, are provided as supplementary information (**Supplementary file, Supplementary note 3**).

## Data and code availability

The data sets and the supplementary movies are available on gitlab (https://gitlab.uliege.be/mipi/published-software/mipi-model-and-simulation-database/-/tree/main/Fitness%20cost%20associated%20with%20cell%20phenotypic%20switching%20drives%20population%20diversification%20dynamics%20and%20controllability). The FlowStoCKS toolbox is on https://gitlab.uliege.be/mipi/published-software/mbms-toolbox/-/tree/main/FlowStoCKS.

## Supplementary information

### Supplementary Information file

**Supplementary movie M1** Time evolution of dotplots FSC-A (Forward Scatter signal) in function of FL1-A (GFP fluorescence signal) obtained based on automated FC for the P_*araB*_:GFP system in *E. coli* cultivated in chemostat

**Supplementary movie M2** Time evolution of dotplots FSC-A (Forward Scatter signal) in function of FL1-A (GFP fluorescence signal) obtained based on automated FC for the P_*lacZ*_:GFP system in *E. coli* cultivated in chemostat

**Supplementary movie M3** Time evolution of dotplots FSC-A (Forward Scatter signal) in function of FL1-A (GFP fluorescence signal) obtained based on automated FC for the P_*bolA*_:GFP system in *E. coli* cultivated in chemostat

**Supplementary movie M4** Time evolution of dotplots FSC-A (Forward Scatter signal) in function of FL1-A (GFP fluorescence signal) obtained based on automated FC for the P_*glc3*_:GFP system in *S. cerevisiae* cultivated in chemostat

**Supplementary movie M5** Time evolution of dotplots FSC-A (Forward Scatter signal) in function of FL1-A (GFP fluorescence signal) obtained based on automated FC for the P_*araB*_:GFP system in *E. coli* cultivated in Segregostat

**Supplementary movie M6** Time evolution of dotplots FSC-A (Forward Scatter signal) in function of FL1-A (GFP fluorescence signal) obtained based on automated FC for the P_*lacZ*_:GFP system in *E. coli* cultivated in Segregostat

**Supplementary movie M7** Time evolution of dotplots FSC-A (Forward Scatter signal) in function of FL1-A (GFP fluorescence signal) obtained based on automated FC for the P_*bolA*_:GFP system in *E. coli* cultivated in Segregostat

**Supplementary movie M8** Time evolution of dotplots FSC-A (Forward Scatter signal) in function of FL1-A (GFP fluorescence signal) obtained based on automated FC for the P_*glc3*_:GFP system in *S. cerevisiae* cultivated in Segregostat

**Supplementary movie M9** Time evolution of dotplots FSC-A (Forward Scatter signal) in function of FL1-A (GFP fluorescence signal) obtained based on automated FC for the P_*spoIIE*_:GFP system in *B. subtilis* cultivated in chemostat (first 40h), followed by Segregostat cultivation.

**Supplementary movie M10** Time evolution of dotplots FSC-A (Forward Scatter signal) in function of FL1-A (GFP fluorescence signal) obtained based on automated FC for the pET28:GFP system in *E. coli* cultivated in chemostat

**Supplementary movie M11** Time evolution of dotplots FSC-A (Forward Scatter signal) in function of FL1-A (GFP fluorescence signal) obtained based on automated FC for the pET28:GFP system in *E. coli* cultivated in Segregostat

**Supplementary movie M12** Cultivation of *S. cerevisiae* carrying a P_*glc3*_:GFP reporter in a MSCC device. Chemically defined medium is constantly perfused into the chamber with a glucose concentration of 0.1mM, leading to a drastic reduction in growth of the colony and full activation of the P_*glc3*_:GFP reporter.

**Supplementary movie M13** Cultivation of *S. cerevisiae* carrying a P_*glc3*_:GFP reporter in a MSCC device. Chemically defined medium is constantly perfused into the chamber with a glucose concentration of 1mM.

**Supplementary movie M14** Cultivation of *S. cerevisiae* carrying a P_*glc3*_:GFP reporter in a dMSCC device. Cultivation conditions are periodically switched between chemically defined media containing glucose at a concentration of 0.1 and 1 mM respectively. The feast-to-famine transitions have been applied with predefined durations i.e., T_0.1mM_ = 0.8 h and T_1mM_ = 3h, for mimicking the Segregostat conditions.

**Supplementary movie M15** FlowStocKS simulations of chemostat cultivations under variable fitness cost.

## Acknowledgement

LH and VV are supported by a FRIA grant provided by the “Fonds de la Recherche Scientifique” FRS-FNRS, from the Walloon region of Belgium. JAM is supported by a post-doctoral grant provided by the Service Public de Wallonie (SPW) and the H2020 program of the European commission (Era-Cobiotech project Contibio). MD is supported by a PhD grant provided by the FRS-FNRS and the H2020 program of the European commission (Era-Net Aquatic Pollutant project ARENA). FD received funding from a research grant provided by the Service Public de Wallonie (SPW) and the H2020 program of the European commission (Era-Cobiotech project ComRaDes).

## Supplementary Information

**Supplementary note 1** Determination of the entropy (H) of the population from automated FC data

Information theory has been used in this work for characterizing the response of cell population to environmental perturbations. This theory involves the computation of entropy H, which can be regarded as a measure of uncertainty about the response of the cell population (output) in function of the environmental stimulation (input). This input-output relationship, which constitute the basis of information theory, will be detailed in **Supplementary note 2**. In this note, we’ll concentrate on the description of the entropy H as a measure of the level of heterogeneity of the population. Entropy can be considered as a measure of uncertainty about the outcome of a draw from a probability distribution^1^ e.g., if we pick randomly a cell in our population, how much are we surprised to pick one cell with a given GFP level? In our case, we can measure the entropy based on fluorescence distribution acquired with automated FC based on the following equation:

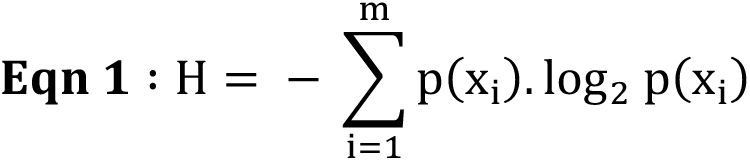

With m being the number of states observed (i.e., GFP classes in our cases) and p being the probability to observe this state. The probability to observe different classes of fluorescence can be easily determined based on automated FC. The computation of H has been exemplified based on fictive population of cells clustered in three fractions according to the level of GFP exhibited by cells (**Figure S1**). As an example, the computation for the first population distribution (**Figure S1A**) is performed as:

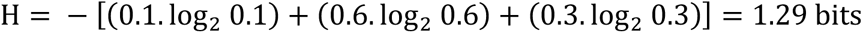

**Figure S1:**
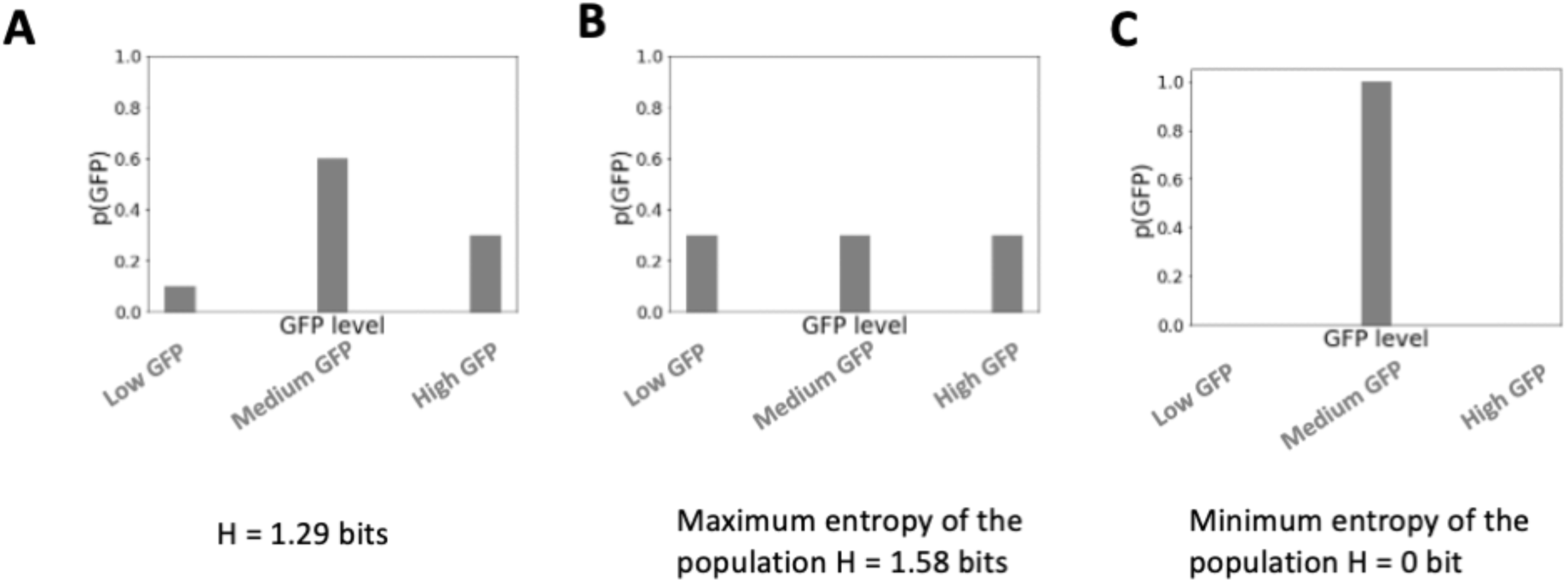
computation of H for cell population clustered in three bins (computation according to equation 1). Each bin corresponds to a subpopulation of cells with a given fluorescence range i.e., low, medium or high.

Based on this first example, the entropy of the population can be either increased (**Figure S1B**), the maximum entropy value being reached when cells are equally distributed into the 3 different clusters. On the opposite, H can be decreased and set to zero when all the cells exhibit the same fluorescence range (**Figure S1C**). This approach has been applied to automated FC data for computing the evolution of H for different types of cell population (**Figure S2** for three out of six of the cell systems investigated). In this case, we applied 50 bins for the computation of H (see Supplementary note 2 for more explanations).

**Figure S2:**
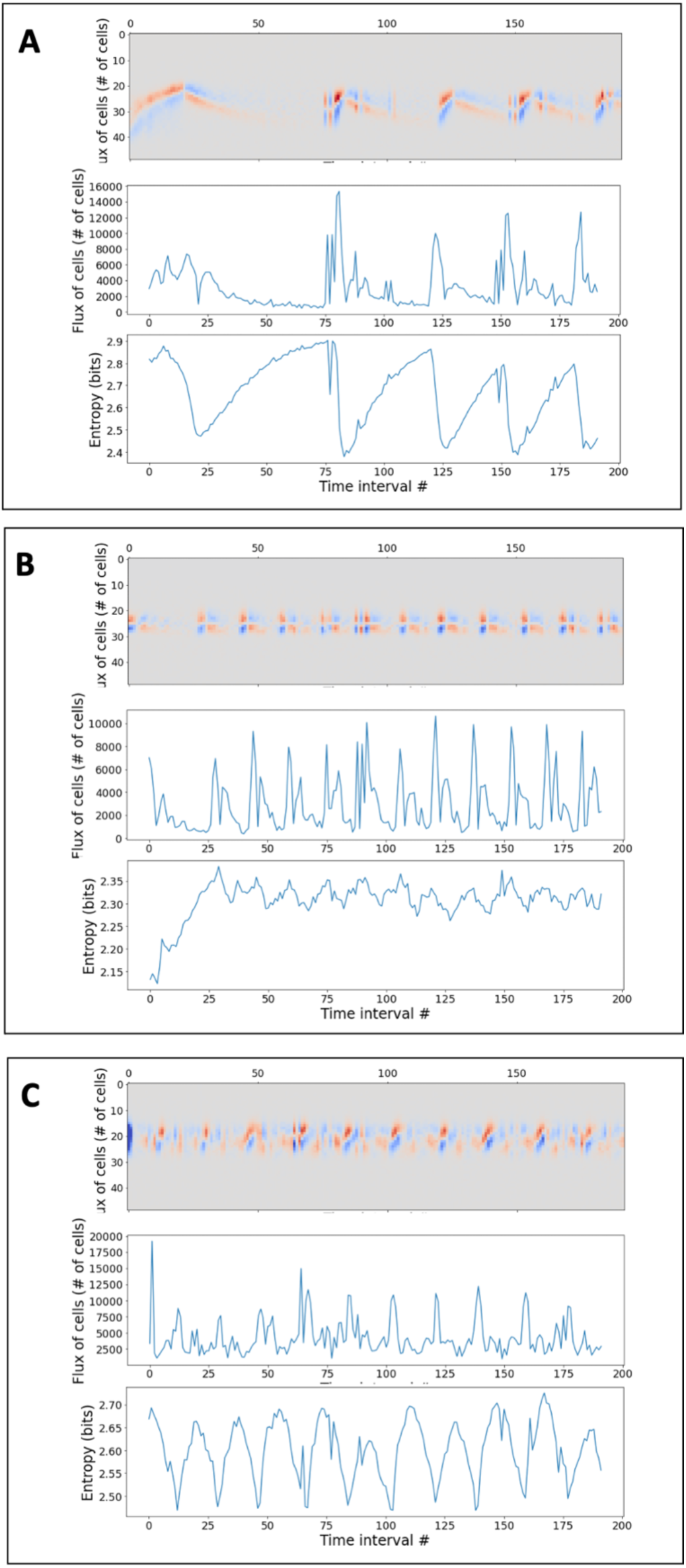
computation of the entropy and flux of cells for the **A** P_*lacZ*_:GFP (*E. coli*), **B** P_*bolA*_:GFP (*E. coli*) and **C** P_*glc3*_:GFP (*S. cerevisiae*) systems.

**Supplementary note 2** Experimental determination of the response function for the P_*araB*_:gfp system in *E. coli* and the P_*glc3*_:gfp system in *S. cerevisiae* and computation of the mutual information (MI)

Information theory relies on the characterization of the input-output relationship for various systems, and has been applied recently to the analysis of signal propagation in biological systems^2,3^. Basically, MI allows to quantify how much we can know about an input (e.g., change in environmental condition) from the output (i.e., in our case, the fluorescence distribution of the population). The first step for the computation of MI is to calculate the entropy of the population exposed to defined environmental conditions i.e., based on the conditional distribution. These conditional distributions represent the response function of our cellular systems and will be characterized in section a and b.

### a. Characterization of the response function for the *E. coli P_araB_:GFP* system

*E. coli* w3110 *ΔaraBAD P_araB_*:GFPmut2 was constructed to characterize the relationship between the inducer concentration and induction profile. The knockout was realized accordingly to Jiang *et al.*^4^ where pTarget was modified with the Fw_sgRNA_20N_Ara primer and the homologous product was constructed from the upstream and downstream fragments generated with Fw_Frag1, Rv_Frag1, FW_Frag2, Rv_Frag2 (**Table S1**).

The deletion of the *araBAD* genes has been done to ensure a perfectly defined concentration of arabinose i.e., no consumption during the trial. From an overnight preculture, 10 flasks (100 ml total volume, 10 ml working volume) with buffered (10 g/l MOPS) mineral salt media were inoculated at an OD_600_ of 0.5. Once the cells were in glucose limited conditions, a solution of arabinose was added to a final concentration ranging from 0 to 2 g/l. Eight concentrations in duplicate were analyzed. Following a delay of 24 minutes, samples analyzed by FC (**Figure S3**) to determine the proportion of cells above the 1000 F.U. threshold. The response function was defined as the ratio of cells above this fluorescence threshold (typical fluorescence value above which a system is considered as induced) over the concentration in arabinose.

**Figure S3:**
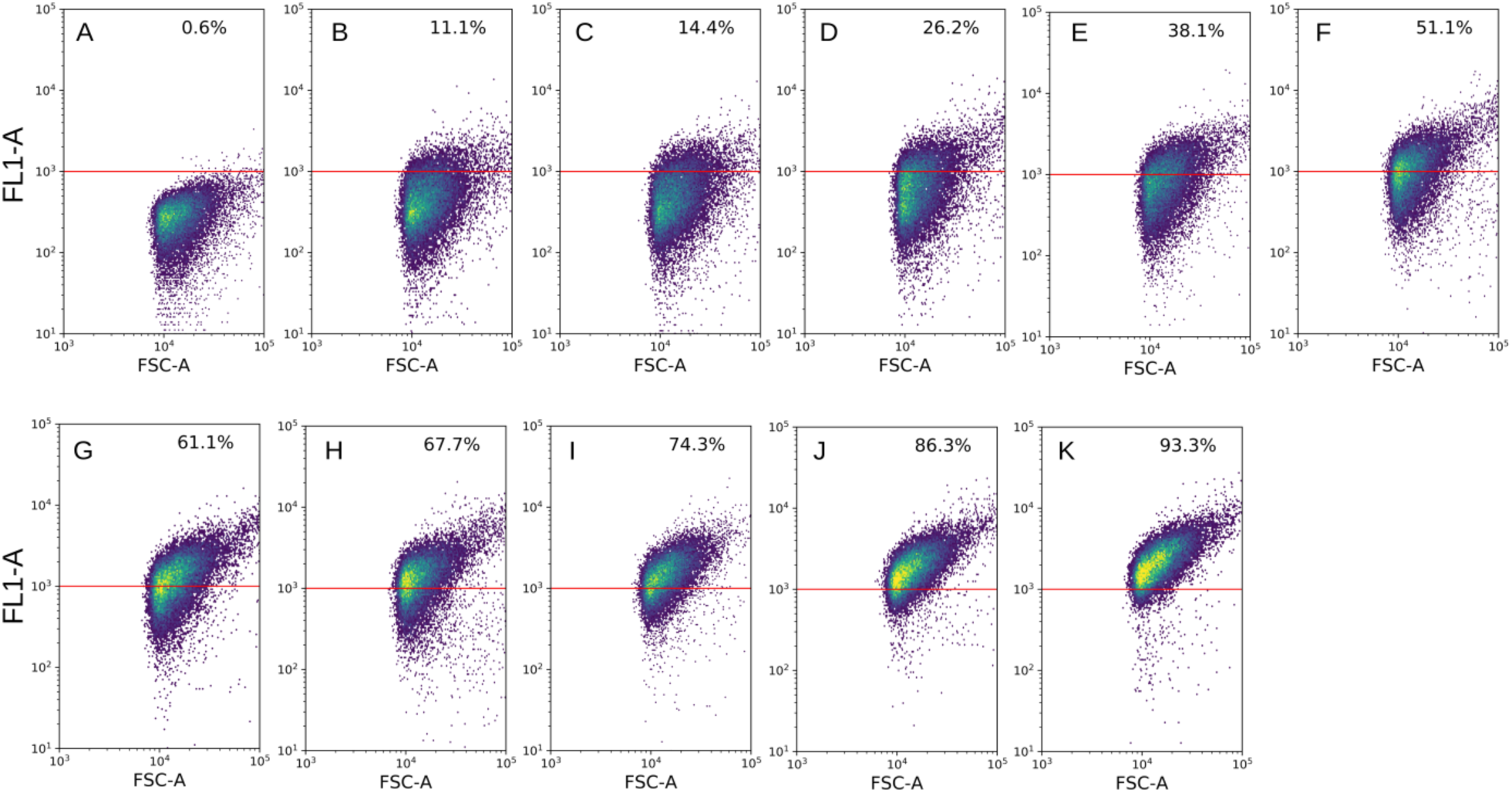
Scatter plots of cell size (FSC-A) versus GFP fluorescence (FL1-A) for *E. coli* w3110 *ΔaraBAD P_araB_*:GFPmut2 exposed to arabinose concentrations (from A to K) of 0, 0.025, 0.05, 0.1, 0.15, 0.20, 0.25, 0.30, 0.50, 1.00 and 2.00 g/L. The redline represents the fluorescence threshold (i.e., 1000 F.U.) used for computing the GFP positive fraction of cells.

**Table S1.**
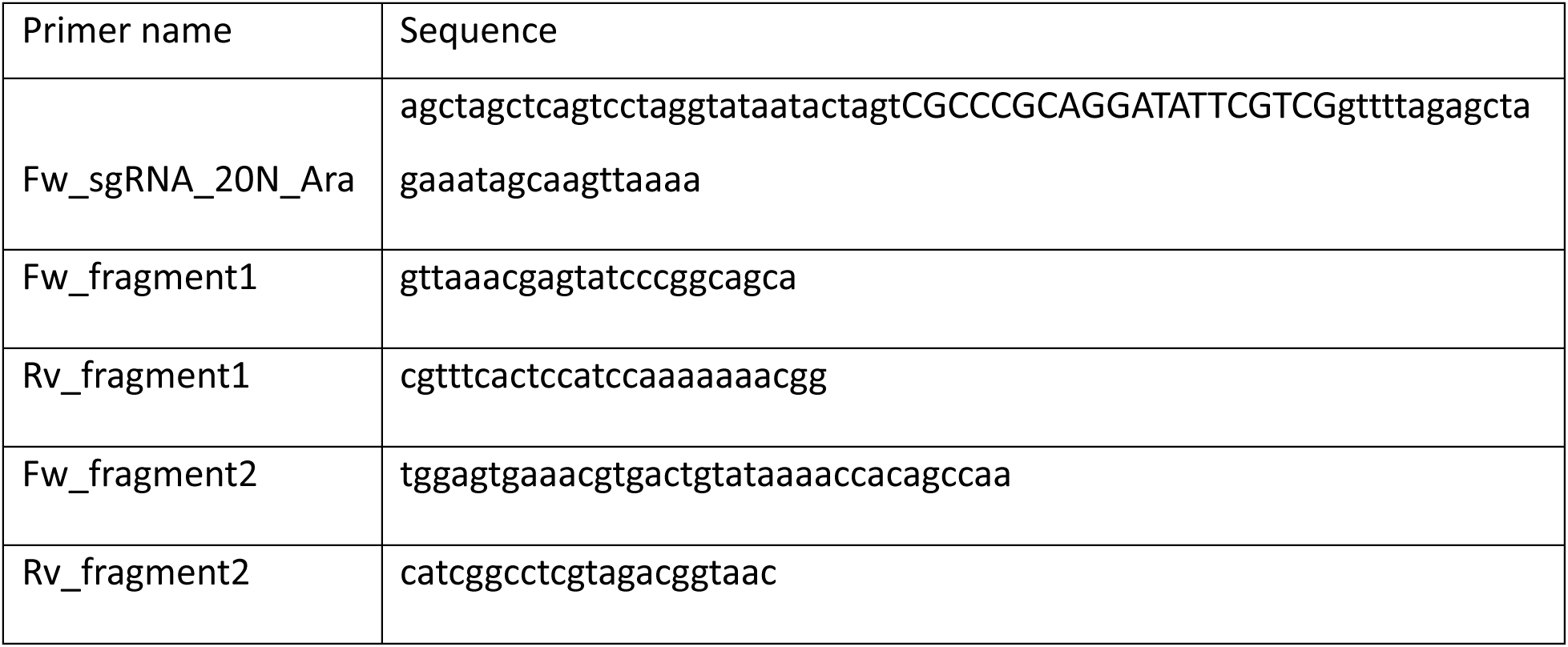

### b. Characterization of the response function for the *S. cerevisiae P_glc3_*:GFP system

The response function of the P_*glc3*_:GFP in *S. cerevisiae* was determined by growing culture at different dilution rates in chemostat (**Figure S4**). For this purpose, the dilution rate of a chemostat was progressively increased in order to release the stress response of the population. This procedure is known as accelerostat (A-stat) and involves the progressive increase of the dilution rate. In our case, the pump flow rate was modified according to step change every 2 hours, resulting in a progressive increase of the dilution rate of 0.002 h^−1^ every hour. This incremental range was chosen in order to ensure pseudo steady-state for each increment. The whole process was followed by automated FC for mapping the GFP distribution of the cell population (**Figure S4**).

**Figure S4:**
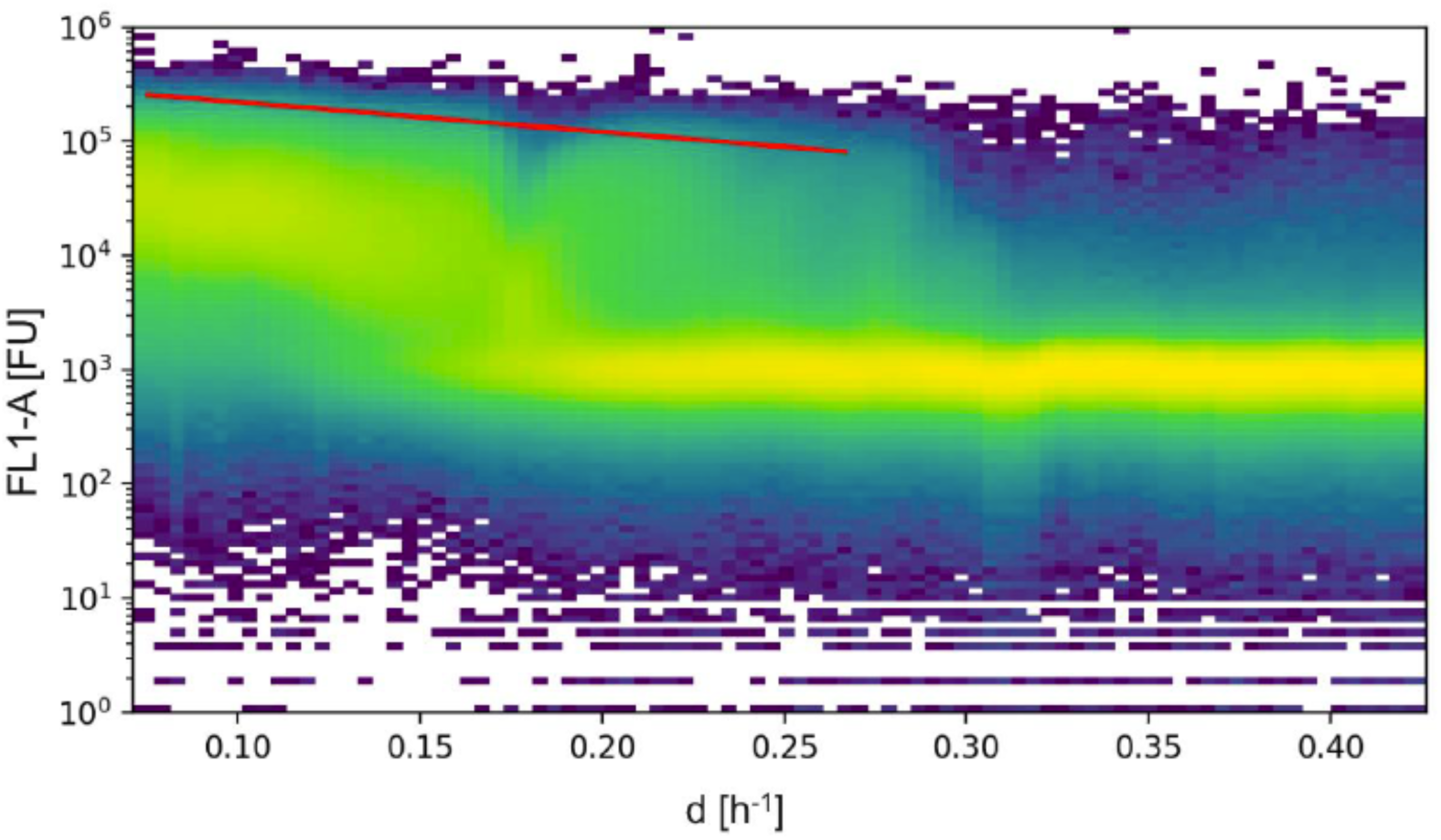
A-stat experiment monitored based on automated FC. The GFP level distribution (FL1-A channel) was determined for each dilution rate (D). The red line highlights the progressive release of the stress response at the population level based on the deactivation of the P_*glc3*_:GFP reporter.

### c. Computation of mutual information (MI) based on the response function of a cell population

Knowing the response function, and the corresponding conditional GFP distribution, it is possible to compute the MI of a specific cellular system. This computation will be exemplified for the P_*glc3*_:GFP reporter in *S. cerevisiae*. The environmental input for this system is the glucose uptake rate determined based on the value of the dilution rate, as well as based on glucose and biomass measurement. The conditional fluorescence distributions were then acquired for different substrate uptake rate and H was computed accordingly (**Figure S5A**). If all the fluorescence distributions are summed up, the corresponding entropy value is the total entropy of the system. MI is then computed according to (**Figure S5B**):

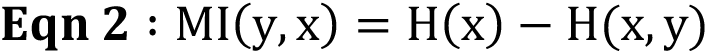

With H(x) being the total entropy for output x and H(x,y) being the conditional entropy computed from the conditional distribution of the output x (x, being GFP distribution).

When doing so, it is important to adjust the number of bins used for computing the entropy. In our case, this number was set to 45 bins and leads to a precise computation of MI without increasing the computing power (**Figure S6**).

**Figure S5:**
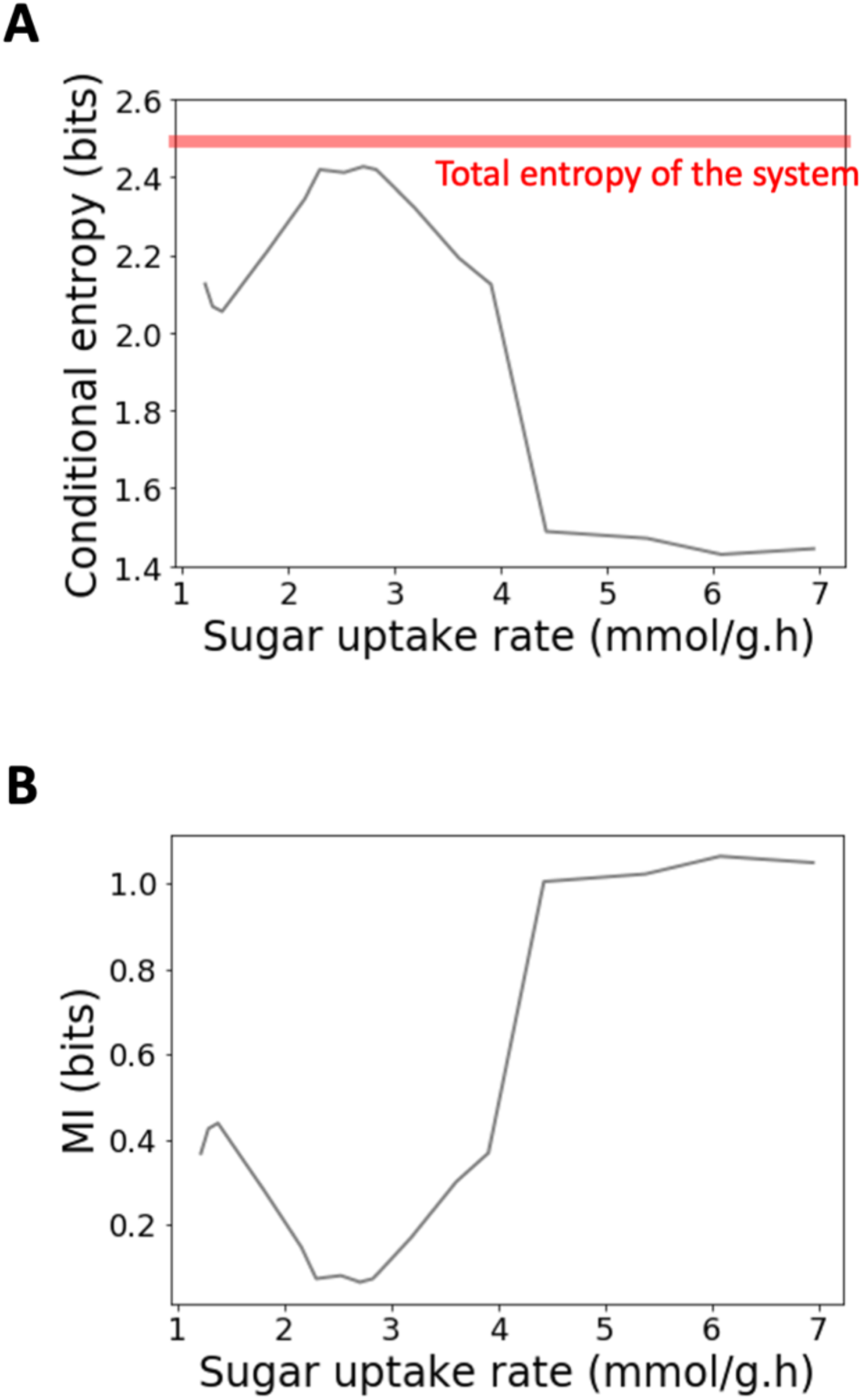
A Evolution of the conditional entropy for the P_*glc3*_:GFP in *S. cerevisiae* exposed to different uptake rates. B MI can be deduced by subtracting the value of the total entropy of the system (2.49 bits in this case) by the corresponding conditional entropy.

**Figure S6:**
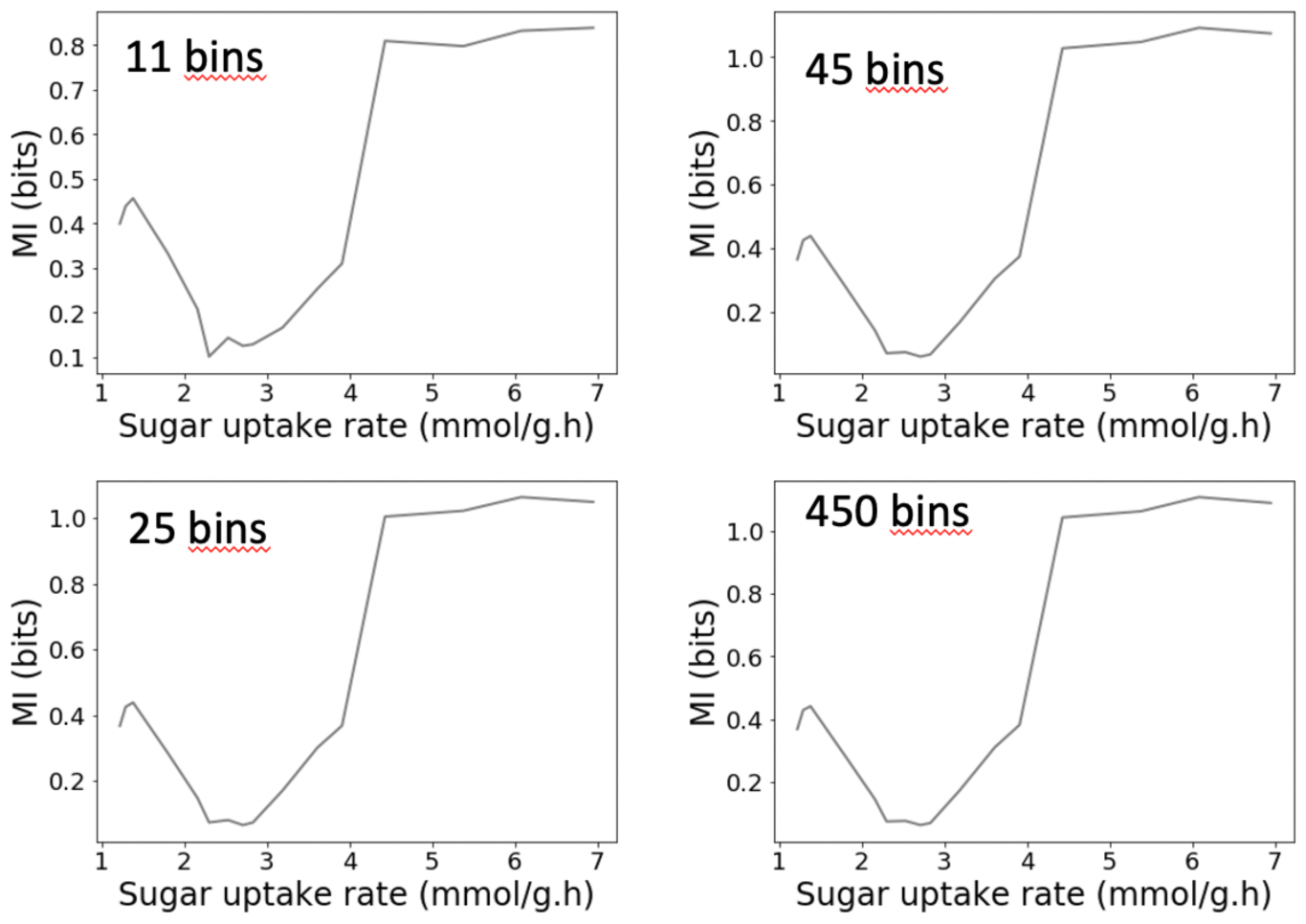
impact of the binning procedure (number of bins considered) on the estimation of MI for the P_*glc3*_:GFP in *S. cerevisiae* (data extracted from A-stat experiments).

**Supplementary note 3** Flow cytometry Stochastic Kinetic Simulator (FlowStocKS)

FlowStocKS is a computational tool designed to simulate the phenotype distributions of a microbial population under different environmental regimes (Chemostat and Segregostat). Following the model framework classification proposed by Hartmann *et al.*^5^, FlowStocKS can be considered as: i) biologically segmented as it considers single cells; ii) abiotically unsegmented as it assumes a homogeneous environment; iii) an unstructured cell model as it does not take intracellular kinetics or metabolic fluxes into consideration. FlowStocKS comprises two modules: the growth module and the switch module. The system is resolved using a Markov chain with discrete time, where the growth module is described by a set of ordinary differential equations (ODE). These ODEs detail how single cells grow (**Eqn 3 and Eqn 4**) and consume (**Eqn 5**) their substrate (S), in accordance with the Monod-type equations that include a non-competitive growth inhibition term.

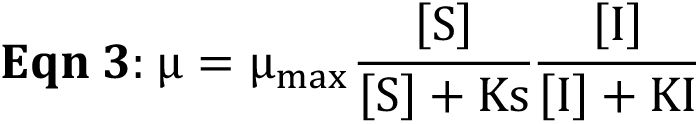

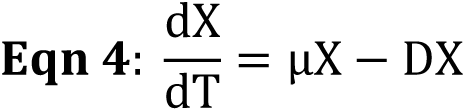

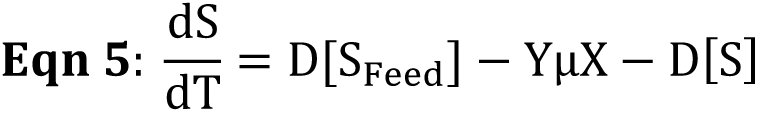

Where:

µ_max_ = Maximal growth rate

µ = Growth rate

*X* = Cell biomass

*KI* = Growth inhibition constant

*K_S_* = Affinity for the substrate

*Y* = Substrate to biomass yield

*D* = Dilution rate

[*S_Feed_*] = Substrate concentration in the feed

[*I*] = Inhibitor concentration

[*S*] = Substrate concentration

In the growth module, single cells are simulated to grow until they double in size, at which point they divide into two daughter cells. Additionally, to simulate continuous cultivations, cells are randomly flushed out of the system based on a probability (*P_out_*) set by the dilution rate and the time step (*T_step_*) used in the simulation (**Eqn 6**).

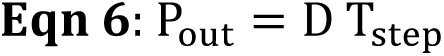

The growth parameters (μ*_max_, KI, Y, K_S_*) are given by the switch module, and define the phenotype of each cell. To initiate the switching process, a cell must first cross a time threshold by accumulating *T_step_*, and this commitment process is governed by a switching probability (P). This probability is determined by a response function, taking the inducer concentration (*i*) as input. The Ρ function is a classical sigmoid function characterized by a steepness (*n*) and a 50% switching probability at concentration (*K*).

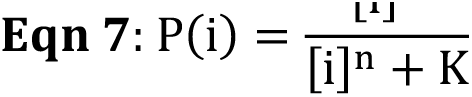

The accumulation of *T_step_* is analogous to the build-up of enzymes that are necessary for the expression of a different phenotype, commonly known in many processes e.g., substrate consumption switching and the diauxic shift time^6,7,8^. Once the time threshold (τ) is reached, the cell switches phenotype. In our experimental set-up, we use GFP-based reporters to track the phenotype switch. Thus, once the time threshold is reached a GFP is produced as a burst following a production rate set by a gamma distribution (*Γ*_(*q_gfp_*,2)_)^9^. Similarly to Taniguchi *et al.*^9^, we assumed that active degradation of GFP or the inhibitor is neglectable and accordingly dilution is determined by cell division.

With this FlowStocKS toolbox, we conducted simulations using a constant time step (*T*_step_) of 1 minute, which is two orders of magnitude faster than the fastest process considered (T_step_<<< µ_max_). In these simulations, we modeled a population of 10,000 cells, each with an initial mass equal to 1/1000th of the initial biomass. Our simulations were based on the growth characteristics of *Saccharomyces cerevisiae* (**Table S1**). The switch to the stressed phenotype is triggered by glucose limitation resulting in growth inhibition. To simplify our analysis, we assumed that the resulting GFP production is linearly correlated with the inhibitor concentration, and thus its concentration was used as the inhibitor

Finally, by varying inhibition value KI (lower value means higher growth inhibition), we were able to compute the phenotype distribution both in Chemostat and Segregostat for different fitness cost associated to switching. All FlowStocKS codes are available on the GitLab page (https://gitlab.uliege.be/mipi/published-software/mbms-toolbox/-/tree/main/FlowStoCKS).

**Table S2:**
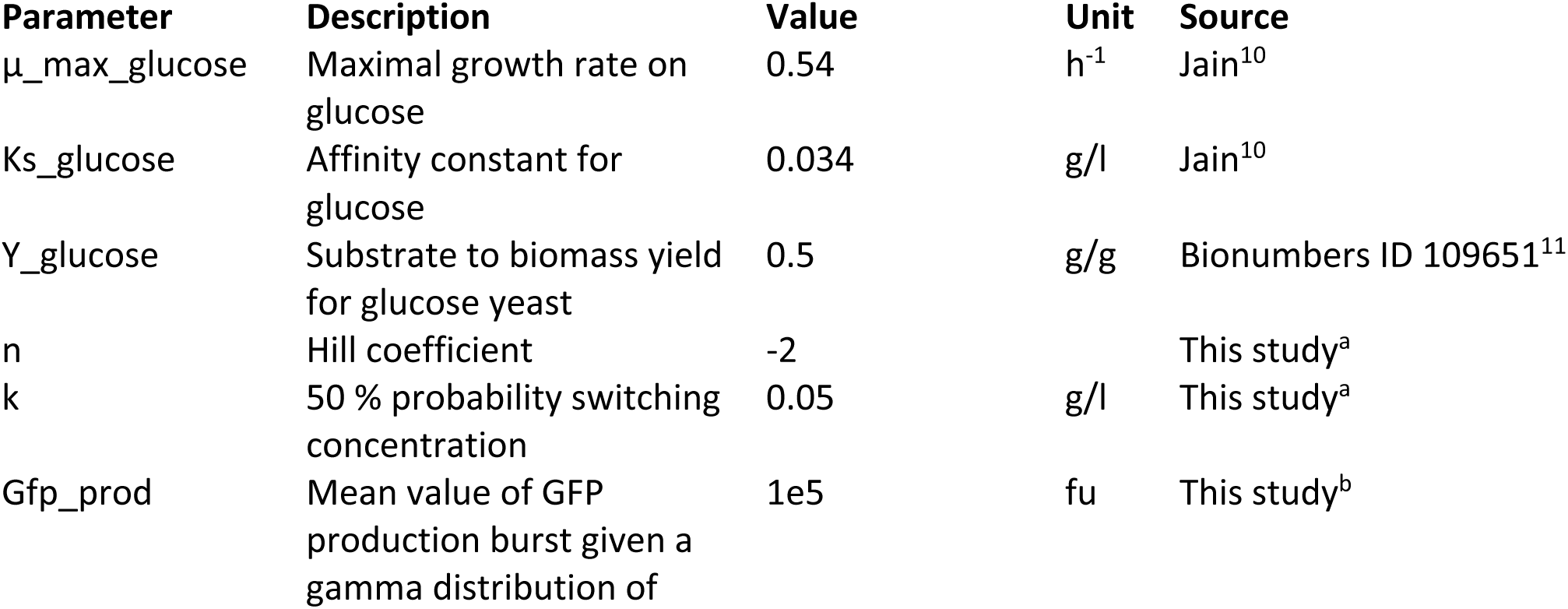
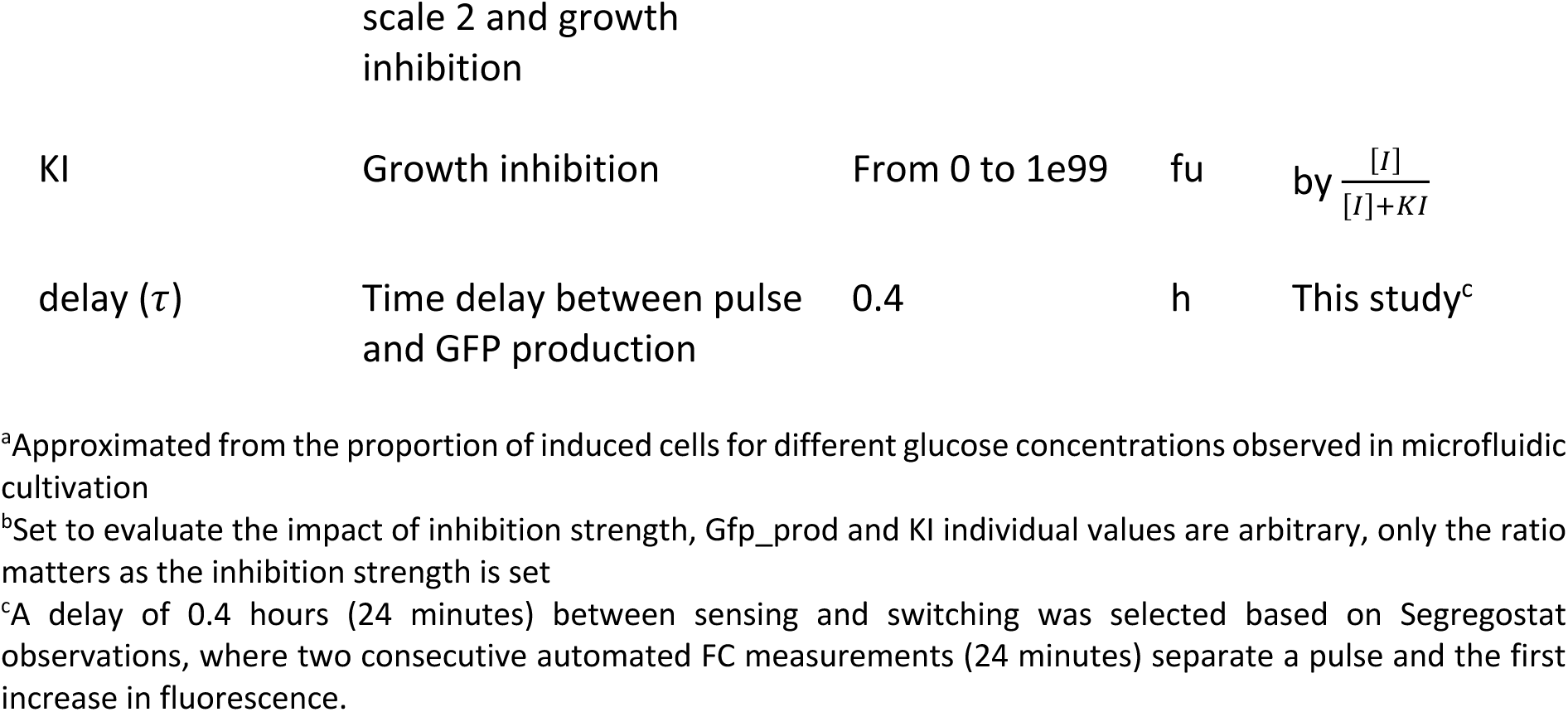
Parameters used for running FlowStocKS simulations

**Supplementary note 4** Operating conditions used for the chemostat and Segregostat experiments

**Table S3.**
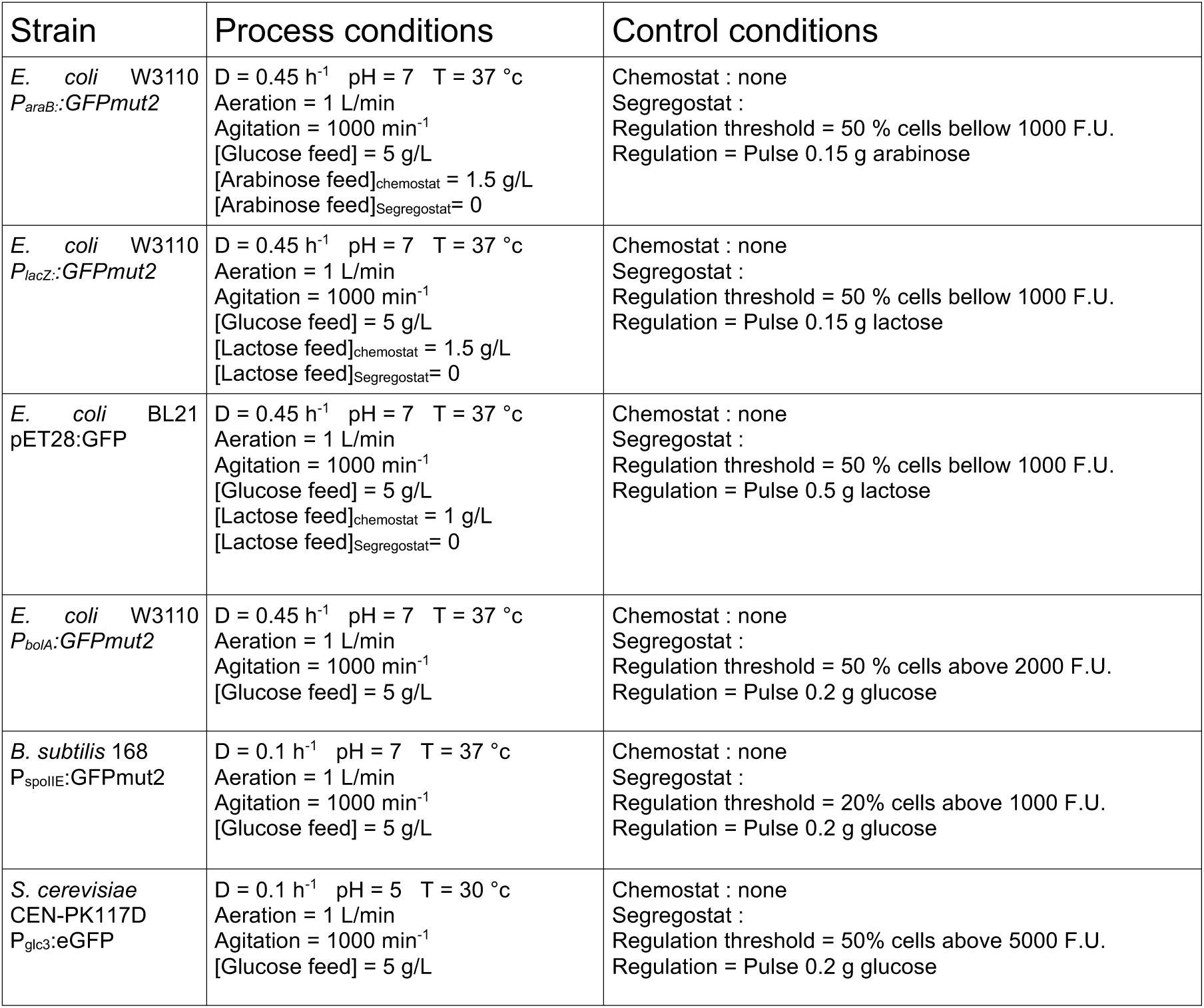

## Notes

### Competing Interest Statement

The authors have declared no competing interest.

